# Live Droplet Single-Cell RNA Sequencing

**DOI:** 10.64898/2026.11.03.763484

**Authors:** David E. Reynolds, Sangmo Lee, Daniel Oh, Shailesh Senthil Kumar, Emily Huynh, Renee-Tyler T. Morales, Sydney Bracht, Phoebe Vallapureddy, Chin Nien Lee, Jianhua Lim, Renis J. Agosto Nieves, Shujing Liu, Gaochen Jin, Xiaowei Xu, Bomyi Lim, Yanxiang Deng, Junhyong Kim, Yoon Ho Roh, Sydney Shaffer, Jina Ko

**Affiliations:** Department of Bioengineering, University of Pennsylvania, Philadelphia, Pennsylvania, USA; Department of Pathology and Laboratory Medicine, University of Pennsylvania, Philadelphia, Pennsylvania, USA; Department of Chemical and Biomolecular Engineering, University of Pennsylvania, Philadelphia, PA, 19104, USA; Graduate Group in Genomics and Computational Biology, Perelman School of Medicine, University of Pennsylvania, Philadelphia, PA, USA; Department of Biology, University of Pennsylvania, Philadelphia, PA, USA; Department of Energy and Chemical Engineering, Incheon National University, 119 Academy-ro, Yeonsu-gu, Incheon 22012, Republic of Korea

**Author notes:** Correspondence: Jina Ko, 422 Curie Blvd, Philadelphia, PA 19104, United States. These authors contributed equally (David E. Reynolds, Sangmo Lee, Daniel Oh).

## Abstract

Single-cell RNA sequencing has emerged as one of the most influential methods of the past decade, enabling the dissection of cellular heterogeneity and the discovery of rare cell types. However, most workflows require cell lysis and provide only terminal snapshots, limiting their ability to capture dynamic cellular trajectories and temporal variations within the same cell. Herein, we present Live Droplet single-cell RNA sequencing (Live D-seq), a technology that enables scalable transcriptome-wide profiling of individual living cells over time. Live D-seq combines longitudinal intracellular sampling of live, barcoded cells with particle-templated emulsification and barcoded hydrogel beads, enabling repeated samples from the same cell lineage to be linked across time points. We showcase Live D-seq’s capabilities by (i) demonstrating minimal cross-contamination in cross-species experiments, (ii) precisely mapping cellular barcodes to hydrogel bead barcodes, and (iii) tracking epithelial-to-mesenchymal transition dynamics in A549 cells over time. Altogether, Live D-seq enables repeated, non-destructive measurements of gene expression in the same single cells, offering a framework to study longitudinal single-cell biology in health and disease.

## 1. Introduction

Single-cell RNA sequencing (scRNA-seq) has transformed our understanding of cellular biology by enabling transcriptome-wide measurements of gene expression within individual cells.^1,2^ Unlike conventional bulk RNA-seq, which averages signals across heterogeneous populations, scRNA-seq maps discrete cellular states and distinct subpopulations directly at single-cell resolution.^3,4^ By revealing cellular heterogeneity and rare cell types, scRNA-seq has been instrumental in charting stem cell differentiation, uncovering localized disease mechanisms, and mapping the immune system’s complex architecture.^5–8^ Despite these advances, most workflows still rely on cell lysis to access intracellular contents. Because these methods are inherently destructive, our ability to track dynamic changes within the same cell over time is limited.^9,10^

Recent technological advances have overcome this limitation by enabling non-destructive RNA measurements in living cells. Live-seq pioneered this approach by using fluidic force microscopy (FluidFM) paired with low-input RNA-seq to capture whole-transcriptome profiles from individual, living cells.^11,12^ Newer adaptations have incorporated electroporation and ion conductance microscopy, which increase overall gene recovery and leverage ion current signals to confirm successful membrane penetration.^13,14^ Beyond physical sampling, other non-destructive strategies rely on chemical probes, metabolic labeling, or CRISPR-based tracking to measure RNA dynamics in real time. For example, photoactivatable systems like transcriptome in vivo analysis (TIVA) and fluorogenic photoaffinity probes enable light-controlled hybridization and tracking of target transcripts, while metabolic incorporation of fluorescent nucleosides allows continuous monitoring of newly synthesized RNA.^15–18^ RNA-targeting CRISPR platforms enable sequence-specific live-cell imaging of endogenous transcription without destroying host cells.^19,20^ While these methods have yielded unprecedented insights into live-cell RNA dynamics, physical sampling methods enable transcriptome-wide coverage with limited throughput, whereas molecular probes offer temporal tracking at the cost of narrow target panels. To bridge this gap, scalable workflows capable of transcriptome-wide profiling of individual living cells are needed.

Here, we present Live Droplet single-cell RNA sequencing (Live D-seq), which enables scalable transcriptome-wide longitudinal profiling of individual living cells in droplets. By coupling particle-templated emulsification (PTE) with transient, mild permeabilization, Live D-seq samples intracellular molecules, including cellular barcodes, from individual living barcoded cells and captures them on magnetic, barcoded hydrogel beads within single droplets. Following sampling and RNA capture, the droplets are demulsified by electro-coalescence to preserve cell viability. The magnetic hydrogel beads are then separated from the cells for next-generation sequencing, while the viable cells are successfully returned to culture for subsequent rounds of longitudinal sampling. To validate the platform, we first demonstrated that cell-to-cell cross-contamination is minimal and comparable to that observed in conventional scRNA-seq. We then confirmed the precise mapping of cellular barcodes to hydrogel bead barcodes, enabling repeated transcriptomic measurements to be linked to the same cell lineages across time points. Finally, we demonstrated longitudinal profiling by tracking epithelial-to-mesenchymal transition (EMT) dynamics in A549 cells. Overall, these results establish Live D-seq as a scalable platform for live scRNA-seq, enabling longitudinal single-cell transcriptomic analysis with broad applications in fundamental biology, diagnostics, and therapeutics.

## 2. Results

### 2.1. Live D-seq workflow

To overcome the limitations of existing single-cell sampling methods, including low scalability, high-cost setups, and high technical delicacy, we developed a robust, scalable approach designed for seamless integration with scRNA-seq workflows. For cellular lineage tracking, we implemented CloneSweeper, a dual-function barcode that enables concurrent live-cell sorting and scRNA-seq (**Fig. 1a**).^21^ Hydrogel beads are uniquely barcoded via a standard split-pool synthesis method (**Fig. 1b**).^22,23^ To quickly and gently co-encapsulate individual barcoded cells and hydrogel beads into single droplets without damaging the cells, we adapted PTE, a vortexing-based strategy previously employed in particle-templated instant partition sequencing (PIP-seq; **Fig. 1c**).^24^ Within these droplets, we perform non-destructive RNA extraction using an optimized hypotonic buffer with a mild permeabilizing agent that creates transient, reversible pores in the cell membrane. This allows intracellular mRNA and cellular barcodes to be released into the extracellular droplet volume, where they are captured on the adjacent barcoded hydrogel bead (**Fig. 1d**). Following RNA capture, droplets are broken, allowing the hydrogel beads to be processed for sequencing while viable cells are returned to culture for downstream tracking (**Fig. 1e**). This gentle intracellular sampling process can be repeated on the same cell population over time. To establish this platform, we systematically optimized each workflow stage, including magnetic hydrogel bead fabrication, cell-hydrogel bead pairing, live-cell RNA extraction, and non-cytotoxic demulsification.

**Figure 1:**
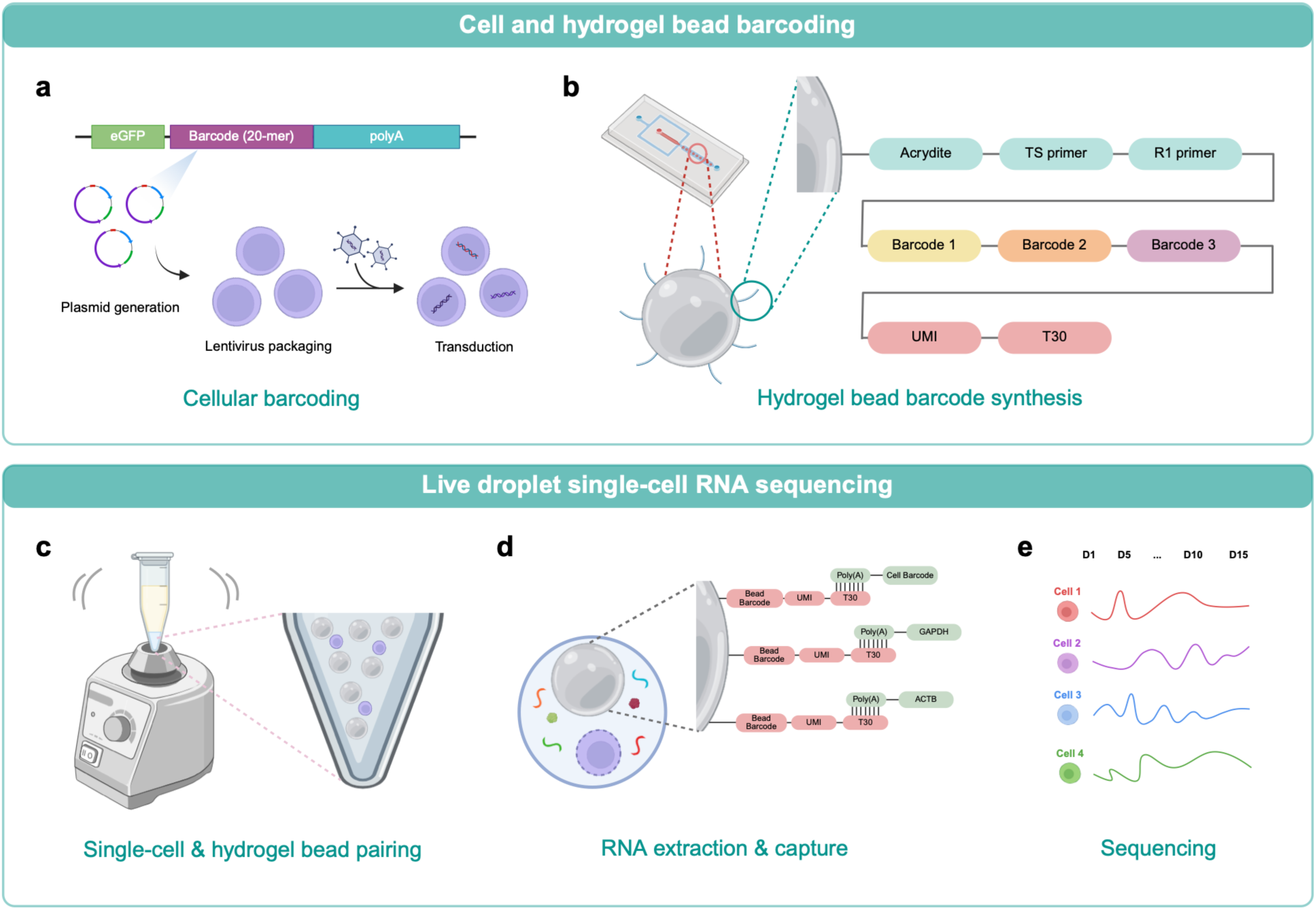
Live D-seq workflow. (**a**) Cellular barcoding with CloneSweeper. Cells are lentivirally transduced with an enhanced green fluorescent protein (eGFP)-barcode-poly(A) construct, giving each cell a heritable, transcribed barcode. (**b**) Magnetic hydrogel beads are generated by droplet microfluidics and barcoded by three-round split-pool synthesis, ending in a unique molecular identifier (UMI) and an oligo(dT)_30_ capture sequence. (**c**) Cells and hydrogel beads are paired in droplets by PTE. (**d**) A hypotonic buffer with a mild permeabilizing agent transiently permeabilizes the cell membrane, releasing mRNA and cellular barcode transcripts for capture on the adjacent hydrogel bead. (**e**) Hydrogel beads are isolated for sequencing while viable cells are returned to culture for repeated sampling. Time points are illustrative. Created with BioRender.com.

### 2.2. Live-cell subsampling preserves viability without perturbing the transcriptome

In traditional droplet-based scRNA-seq, cells are lysed within droplets and only the barcoded transcripts are recovered for sequencing. In Live D-seq, however, cell viability is essential: both intact cells and hydrogel beads must be recovered—the cells for downstream culture and the hydrogel beads for sequencing. Because cells and hydrogel beads are both suspended in the same aqueous solution, physical separation without damaging the cells is challenging. To overcome this, we incorporated magnetic nanoparticles into the hydrogel beads, allowing the hydrogel beads to be magnetically retained while the cells remain in suspension for intact recovery. Following established protocols, we synthesized magnetic poly(ethylene glycol) diacrylate (PEGDA) hydrogel beads using droplet microfluidics.^25^ We encapsulated magnetic nanoparticles within hydrogel precursor droplets before photopolymerization and subsequent droplet breakage and washing yielded freestanding magnetic hydrogel beads (**Fig. 2a**).

**Figure 2:**
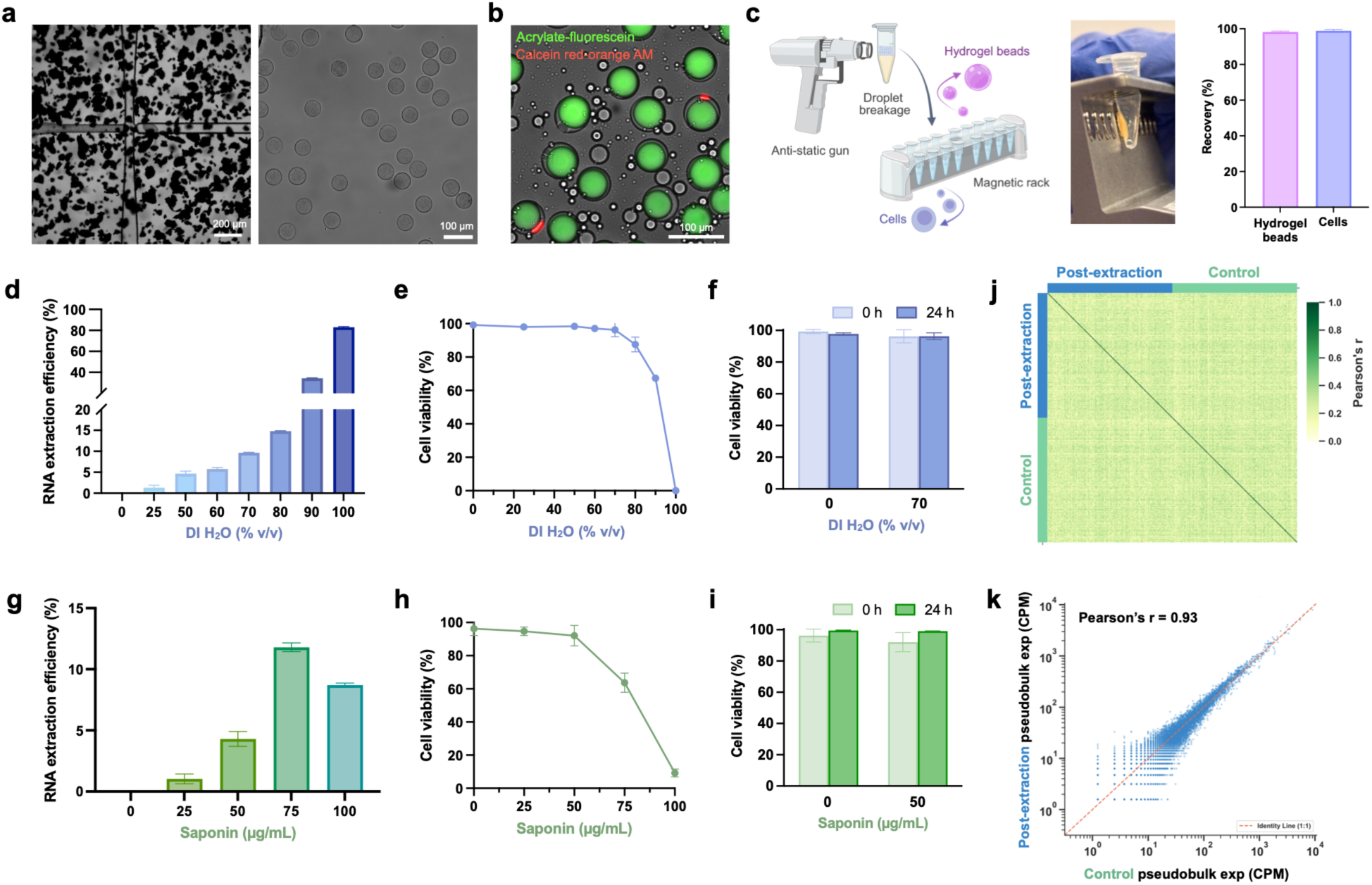
Live-cell subsampling preserves viability without perturbing the transcriptome. (**a**) Fabrication of magnetic hydrogel beads by droplet microfluidics (left) and the resulting freestanding hydrogel beads after droplet breakage and washing (right). (**b**) Single-cell and hydrogel bead pairing by PTE. Hydrogel beads are labeled with acrylate-fluorescein (green) and cells with calcein red-orange AM (red). (**c**) Demulsification with an anti-static gun and magnetic separation of hydrogel beads from cells, with recovery of hydrogel beads and cells. Created with BioRender.com. (**d–f**) RNA extraction efficiency (**d**), immediate cell viability (**e**), and viability at 0 and 24 h (**f**) of HEK293T cells across DI water concentrations, without saponin. Extraction efficiency in (**d**) is shown relative to 0% (v/v) DI water. (**g–i**) Additional RNA extraction efficiency (**g**), immediate cell viability (**h**), and viability at 0 and 24 h (**i**) across saponin concentrations in 70% (v/v) DI water. Extraction efficiency in (**g**) is shown relative to 0 µg/mL saponin. (**j**) Pairwise cell-cell Pearson correlation of HEK293T cells 48 h after Live D-seq compared with untreated controls (150 highest-UMI cells per group). (**k**) Pseudobulk expression of all post-extraction versus control cells (*n* = 260 post-extraction and 271 control cells). Data in (**c–i**) are shown as mean ± s.d. (*n* = 3 technical replicates). Scale bars: 200 µm and 100 µm (**a**), 100 µm (**b**).

With hydrogel bead synthesis established, we next evaluated whether individual hydrogel beads could be efficiently paired with single cells. Our approach utilizes PTE, in which a small volume (<10 µL) of a cell-hydrogel bead aqueous mixture is vortexed in a larger volume of oil (500 µL). To optimize single-cell and single-hydrogel bead pairing, we varied the cell-to-hydrogel bead ratio. Increasing the proportion of cells relative to a fixed number of hydrogel beads increased the fraction of hydrogel beads paired with exactly one cell from ∼5% at a 5% ratio to ∼7% at 10% and ∼10% at 20% (**Supplementary Fig. 1**). However, increasing the cell-to-hydrogel bead ratio beyond 10% (up to 20%) resulted in more droplets containing a single hydrogel bead and multiple cells. We therefore selected a 10% cell-to-hydrogel bead ratio for subsequent experiments (**Fig. 2b**). Following successful pairing, we sought an effective method to demulsify the droplets without compromising cell viability. While chemical demulsifiers such as 1H,1H,2H,2H-perfluoro-1-octanol (PFO) are widely used, they can exert cytotoxic effects. To circumvent this, we implemented a Zerostat anti-static gun to induce electro-coalescence. This approach brings droplets into close contact, displaces the intervening oil layer, and disrupts the surfactant interface without compromising cell viability.^26^ After approximately 30 s of actuation with the anti-static gun, the emulsion separated into distinct aqueous and oil phases. The aqueous phase was collected and transferred to a tube on a magnetic rack to isolate the magnetic hydrogel beads from the supernatant. The recovered cells were then returned to culture while the hydrogel beads were prepared for sequencing, achieving >95% recovery of both cells and hydrogel beads (**Fig. 2c**).

With validated cell-hydrogel bead pairing and recovery, we optimized the protocol for live-cell RNA extraction. To enable rapid, reversible extraction while minimizing cellular perturbation and remaining compatible with PTE, we employed a combination of deionized (DI) water and saponin. In this process, DI water creates a hypotonic environment that induces osmotic swelling and increases membrane tension, while saponin permeabilizes the plasma membrane by interacting with cholesterol and forming transient pores.^27–30^ Because cholesterol is abundant in the plasma membrane but scarce in intracellular membranes, low saponin concentrations permeabilize the plasma membrane while leaving organelle membranes largely intact.^31–33^ Cytosolic mRNA can therefore diffuse out while the nucleus and mitochondria are retained, and the small pores formed under these conditions allow the membrane to reseal after extraction. This allows intracellular molecules to be released into the extracellular space for hydrogel bead capture. Crucially, buffer ratios must be carefully calibrated to prevent cell over-swelling or mechanical rupture of the plasma membrane. While hypotonic treatment has previously been used for intracellular delivery, we repurposed it for non-destructive, live-cell RNA extraction. To determine the conditions for RNA extraction, we initially varied the ratio of DI water to 1% bovine serum albumin (BSA) in Dulbecco’s phosphate-buffered saline (DPBS) without calcium and magnesium, where BSA serves as a neutral buffer to maintain high cell viability, using HEK293T cells in the absence of hydrogel beads and oil (30 min incubation at room temperature). As expected, increasing the proportion of DI water led to a corresponding increase in RNA yield per cell (**Fig. 2d**). However, cell viability dropped below 90% immediately after treatment with 80% (v/v) DI water, and complete cell lysis occurred at 90–100% (v/v) (**Fig. 2e**). Because 70% (v/v) DI water maintained high cell viability (∼96%) while yielding an RNA extraction efficiency of ∼10% per cell, we selected this condition for subsequent experiments. An extraction efficiency of 5–15% is ideal for live-cell RNA profiling, as prior studies have shown this range provides sufficient mRNA for sequencing while preserving cell viability.^11,12,34^ To confirm long-term survival, we evaluated cell viability 24 h post-extraction, which remained above 95% (**Fig. 2f**). To assess generalizability, we further evaluated RNA extraction efficiency and cell viability in NIH-3T3, A549, Jurkat, and primary T cells under the same hypotonic buffer conditions (**Extended Data Fig. 1a–c**).

To further improve RNA yield, we evaluated saponin as a mild membrane-permeabilizing agent. Testing four concentrations showed that saponin increased RNA yield by an additional 1– 12% per cell beyond 70% (v/v) DI water alone, with the increase plateauing at 8–12% at 75–100 µg/mL (**Fig. 2g**). However, cell viability measured immediately post-treatment dropped sharply below 70% at these higher concentrations (**Fig. 2h**). Consequently, we selected 50 µg/mL saponin, which added ∼4% (∼14% total extraction efficiency per cell) while maintaining cell viability above 95% at 24 h post-extraction (**Fig. 2i**). To visualize molecular leakage during extraction, we imaged calcein green AM–loaded HEK293T cells over 30 min in saponin buffer (50 µg/mL) with 0%, 70%, or 100% (v/v) DI water. At 70% (v/v) DI water, cells swelled and gradually leaked calcein but stayed mostly viable, whereas 100% (v/v) DI water caused rapid signal loss and complete lysis (**Supplementary Fig. 2a,b**). Comparison of treated (70% (v/v) DI water + 50 µg/mL saponin) and untreated HEK293T cells 24 h post-extraction, along with cell surface staining for membrane proteins like CD9, confirmed a healthy phenotype and continued cell proliferation (**Supplementary Fig. 3a,b**). To assess broader applicability, we validated these buffer conditions across NIH-3T3, A549, Jurkat, and primary T cells, confirming high viability (**Extended Data Fig. 1d–f**) and healthy, proliferating cells at 0 and 24 h post-treatment (**Extended Data Fig. 1g–j**). These optimization steps established the standard extraction buffer for subsequent experiments, while highlighting the potential need for further adjustments across cell types and culture conditions due to differences in membrane sensitivity.

With our live single-cell subsampling workflow optimized and extraction buffer conditions established in bulk, we next verified that droplet-based extraction does not perturb baseline cellular transcriptomes. We performed Live D-seq on HEK293T cells using magnetic hydrogel beads suspended in our hypotonic buffer (70% (v/v) DI water and 50 µg/mL saponin) with a 25 min droplet incubation. Following demulsification, recovered cells were returned to culture for 48 h and subsequently analyzed by conventional droplet-based scRNA-seq alongside untreated controls. To assess whether extraction altered cellular transcriptional states, we first examined pairwise cell-cell correlations. The correlation matrix showed no distinct separation between post-extraction and control cells, with between-group correlations (mean *r* = 0.25) comparable to within-group correlations (mean *r* = 0.25 and 0.26 for post-extraction and control cells, respectively), demonstrating minimal transcriptomic perturbation (**Fig. 2j**). To further assess the preservation of overall gene expression profiles, we compared pseudobulk expression between the two populations. This analysis between post-extraction and control cells yielded a strong correlation (Pearson’s *r* = 0.93), confirming minimal perturbation following live-cell extraction (**Fig. 2k**). Collectively, these findings establish a scalable framework for live single-cell subsampling that enables parallel processing of cells in droplets, while achieving efficient RNA extraction, high cell viability, and minimal transcriptomic perturbation.

### 2.3. Live D-seq achieves single-cell resolution with minimal cross-talk

To demonstrate that Live D-seq achieves true single-cell resolution, we performed a series of benchmarking experiments. To uniquely index extracted transcripts and minimize cross-talk, we incorporated combinatorially barcoded hydrogel beads. Using a three-round split-pool synthesis strategy, we functionalized magnetic hydrogel beads with three barcode sequences followed by a terminal oligo(dT)_30_ sequence for mRNA capture (**Supplementary Table 1**). Successful barcode functionalization was confirmed by probing hydrogel beads with a FAM-labeled poly(A) oligonucleotide. Fluorescence microscopy demonstrated strong, uniform signal intensity across the barcoded magnetic hydrogel beads, with negligible background in non-barcoded controls (**Supplementary Fig. 4a**). To confirm that our barcoded magnetic hydrogel beads effectively capture and process mRNA, we incubated them with soluble total RNA. Following reverse transcription (RT), PCR amplification, and solid-phase reversible immobilization (SPRI) bead purification, TapeStation analysis revealed a broad cDNA size distribution characteristic of full-length mRNA recovery (**Supplementary Fig. 4b**).

We next performed a cross-species experiment to assess single-cell resolution and potential cross-talk. Human (HEK293T) and mouse (NIH-3T3) cells were co-encapsulated with barcoded magnetic hydrogel beads in hypotonic buffer (70% (v/v) DI water containing 50 µg/mL saponin). Following droplet breakage and magnetic hydrogel bead-cell separation, the mRNA-bound hydrogel beads were processed for sequencing. As a reference, a parallel sample was processed using conventional droplet-based scRNA-seq with in-droplet lysis on the same platform. Based on the barnyard plots, both scRNA-seq and Live D-seq yielded distinct human and mouse cell populations. The doublet rates remained low across both platforms at 1.6% for scRNA-seq and 4.6% for Live D-seq. Cell capture was comparable between methods (217–248 cells captured per run from ∼1,200 input cells), demonstrating comparable performance whether cells were lysed or sampled live (**Fig. 3a**). Both HEK293T and NIH-3T3 cells exhibited normal cell spreading and remained highly viable 24 h after Live D-seq (**Supplementary Fig. 5**).

**Figure 3:**
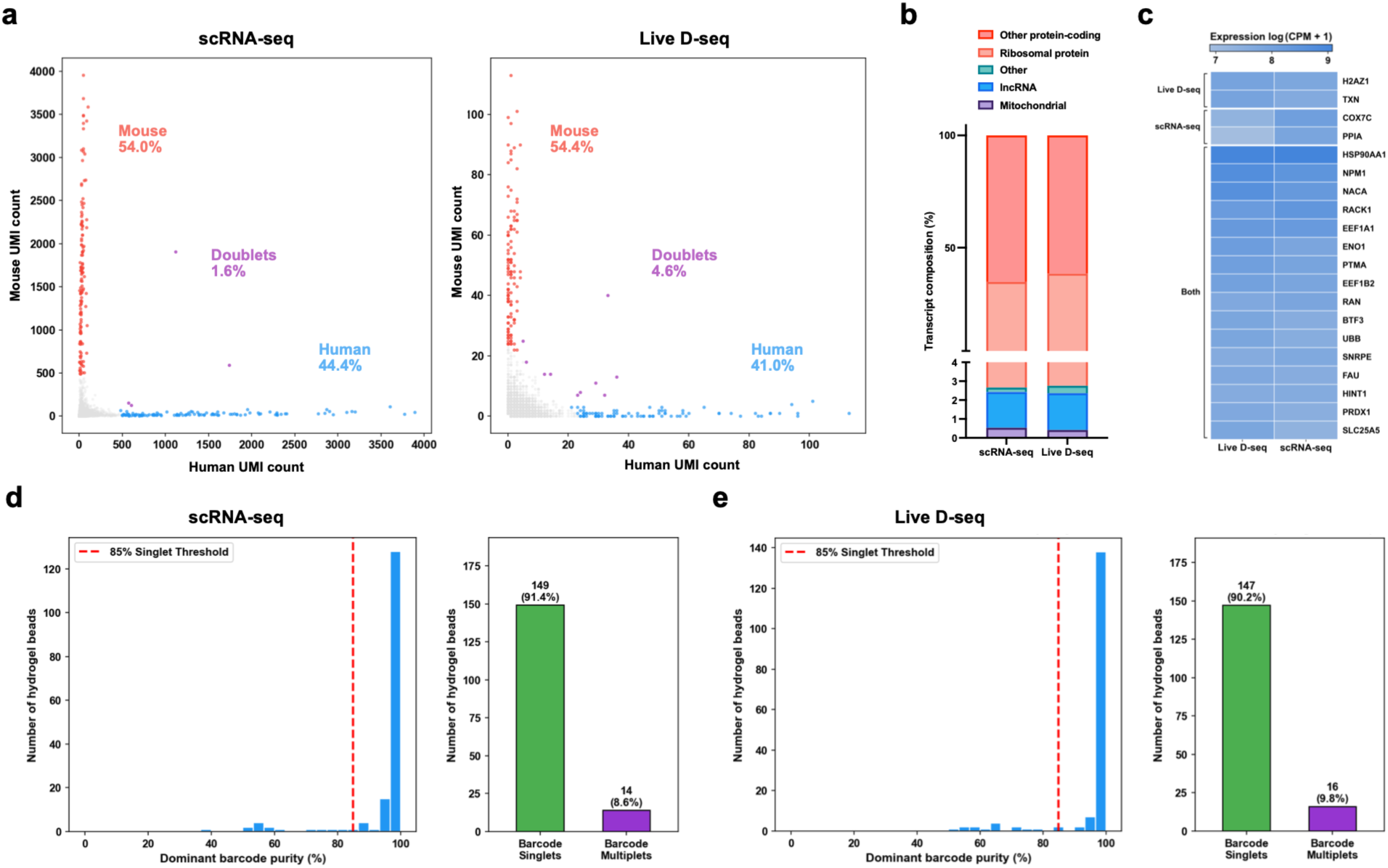
Live D-seq achieves single-cell resolution with minimal cross-talk. (**a**) Barnyard plots of human (HEK293T) and mouse (NIH-3T3) cells profiled by single-cell RNA sequencing (scRNA-seq; left) and Live D-seq (right). Each point is one hydrogel bead, colored by species assignment: human (blue), mouse (red), or doublet (purple). Hydrogel beads were assigned to a species when at least 85% of their UMIs mapped to that species. Only hydrogel beads above 500 UMIs (scRNA-seq) or 24 UMIs (Live D-seq) were classified and included in (**b**) and (**c**). (**b**) Transcript composition of human cells by gene category, calculated using all genes. (**c**) Expression of the most abundant genes in human cells, excluding ribosomal protein genes, grouped by whether they were among the most abundant in Live D-seq only, scRNA-seq only, or both. (**d,e**) Cellular barcode purity on hydrogel beads for barcoded A549 cells profiled by scRNA-seq (**d**) and Live D-seq (**e**), for hydrogel beads whose dominant cellular barcode had at least 100 (scRNA-seq) or 15 (Live D-seq) read pairs. Histograms show the fraction of cellular barcode reads on each hydrogel bead assigned to its dominant barcode. Hydrogel beads at or above the 85% threshold (dashed line) were classified as barcode singlets. Bar plots show the numbers of barcode singlets and multiplets.

To check that Live D-seq captures transcriptomes that resemble those from conventional scRNA-seq, we compared expression and library metrics between the two methods. Sequencing saturation, read depth, and uniquely mapped reads for both libraries are shown in **Extended Data Fig. 2a–c**. As expected, Live D-seq recovered fewer UMIs and genes per cell than scRNA-seq in both human and mouse cells (**Extended Data Fig. 2d**). Even so, its transcriptomes closely matched scRNA-seq. Transcript composition was nearly the same in both species: other protein-coding genes made up most of each library ∼60–65%, ribosomal protein transcripts ∼30–35%, lncRNAs ∼2%, and mitochondrial transcripts <1% (**Fig. 3b** and **Extended Data Fig. 2e**). Mitochondrial UMI fractions were low in both methods (<2%), suggesting Live D-seq sampling does not cause transcriptional signs of cell damage (**Extended Data Fig. 2f**).^35^ As expected, almost every gene detected by Live D-seq was also detected by scRNA-seq (1,584 of 1,662 human genes, 95%; 2,098 of 2,170 mouse genes, 97%; **Extended Data Fig. 2g**). In scRNA-seq, these shared genes averaged far more UMIs per gene than genes seen only by scRNA-seq (∼68 vs. ∼5 in human; ∼78 vs. ∼6 in mouse), indicating that Live D-seq primarily misses low-abundance transcripts. Despite these differences, pseudobulk expression remained moderately correlated between the two methods (Pearson’s *r* = 0.61 for human and 0.65 for mouse, **Extended Data Fig. 2h**). The most abundant genes largely overlapped between the two methods. Excluding ribosomal protein genes, shared genes included *HSP90AA1*, *NPM1*, *EEF1A1*, *RACK1*, and *ENO1* in human and *Fth1*, *Vim*, *Tpt1*, *Lgals1*, and *Hsp90ab1* in mouse. This overlap held when ribosomal protein genes were included (**Fig. 3c** and **Extended Data Fig. 2i–k**), with some minor differences in the most abundant genes detected in each method (e.g., *H2AZ1* and *TXN* in Live D-seq versus *COX7C* and *PPIA* in scRNA-seq for human).

After demonstrating single-cell resolution, we next applied Live D-seq to A549 cells labeled with unique cellular barcodes (CloneSweeper). To validate the CloneSweeper system before applying Live D-seq, eGFP-positive barcoded A549 cells were isolated by fluorescence-activated cell sorting (FACS) and expanded in culture. We first confirmed barcode expression in bulk total RNA, observing a 2.2 × 10^6^-fold enrichment of the cellular barcode in barcoded versus non-barcoded cells, while housekeeping genes (*ACTB* and *GAPDH*) remained unchanged (∼1–2-fold change; **Supplementary Fig. 4c**). Next, to confirm functional capture on our platform, total RNA was co-incubated with barcoded magnetic hydrogel beads. Subsequent RT-qPCR demonstrated robust, specific on-hydrogel bead capture, yielding a ∼1.6 × 10^4^-fold enrichment of the cellular barcode with minimal variation in housekeeping controls (**Supplementary Fig. 4d**). We then co-encapsulated barcoded A549 cells with barcoded magnetic hydrogel beads in hypotonic buffer (∼70% (v/v) DI water). We omitted saponin because our optimization steps showed it provided negligible improvement in RNA extraction from A549 cells. After sequencing, we identified the cellular barcodes captured on each hydrogel bead and calculated barcode purity as the fraction of reads assigned to the dominant cellular barcode. Hydrogel beads with >85% barcode purity were classified as singlets, with most hydrogel beads in both datasets exhibiting nearly 100% purity (**Fig. 3d,e**). Live D-seq classified 90.2% of hydrogel beads as barcode singlets and 9.8% as multiplets. scRNA-seq of the same barcoded cells classified 91.4% of hydrogel beads as singlets and 8.6% as multiplets. Together, these results show that Live D-seq enables accurate cellular barcode assignment at single-cell resolution, with performance comparable to conventional scRNA-seq.

### 2.4. Live D-seq tracks EMT dynamics in single cells

To demonstrate the capability of Live D-seq for longitudinal single-cell profiling, we tracked EMT dynamics in A549 cells following TGF-β stimulation (**Fig. 4a**). The same population of barcoded cells was sampled by Live D-seq one day before TGF-β treatment (D-1) and one day after treatment (D1), then fully lysed on Day 3 (D3). Because cells may divide between time points, longitudinal tracking was performed at the cell-lineage level using heritable cellular barcodes. For scRNA-seq controls, cells from separate wells were lysed at each time point. Before the time-course experiment, we confirmed by bright-field microscopy that treatment with 4 ng/mL TGF-β, a previously established working concentration, induced EMT-associated morphological changes.^36,37^ By D1, cells had become elongated and spindle-shaped, consistent with a mesenchymal state (**Fig. 4b**). Imaging throughout the experiment showed that cells sampled by Live D-seq remained eGFP-positive and exhibited progressively elongated, spindle-shaped morphologies consistent with EMT. Because only cells recovered after sampling were returned to culture, these cultures had lower cell densities than the unsampled controls, which continued to proliferate to confluence (**Supplementary Fig. 6a,b**). To confirm that the same cell lineages were being tracked, we compared the cellular barcodes recovered at D-1 and D1. For each cell lineage, the D-1 barcode had a clear counterpart at D1, readily distinguished from all other barcodes by its low Hamming distance (**Fig. 4c**). This allowed us to follow these lineages across both time points.

**Figure 4:**
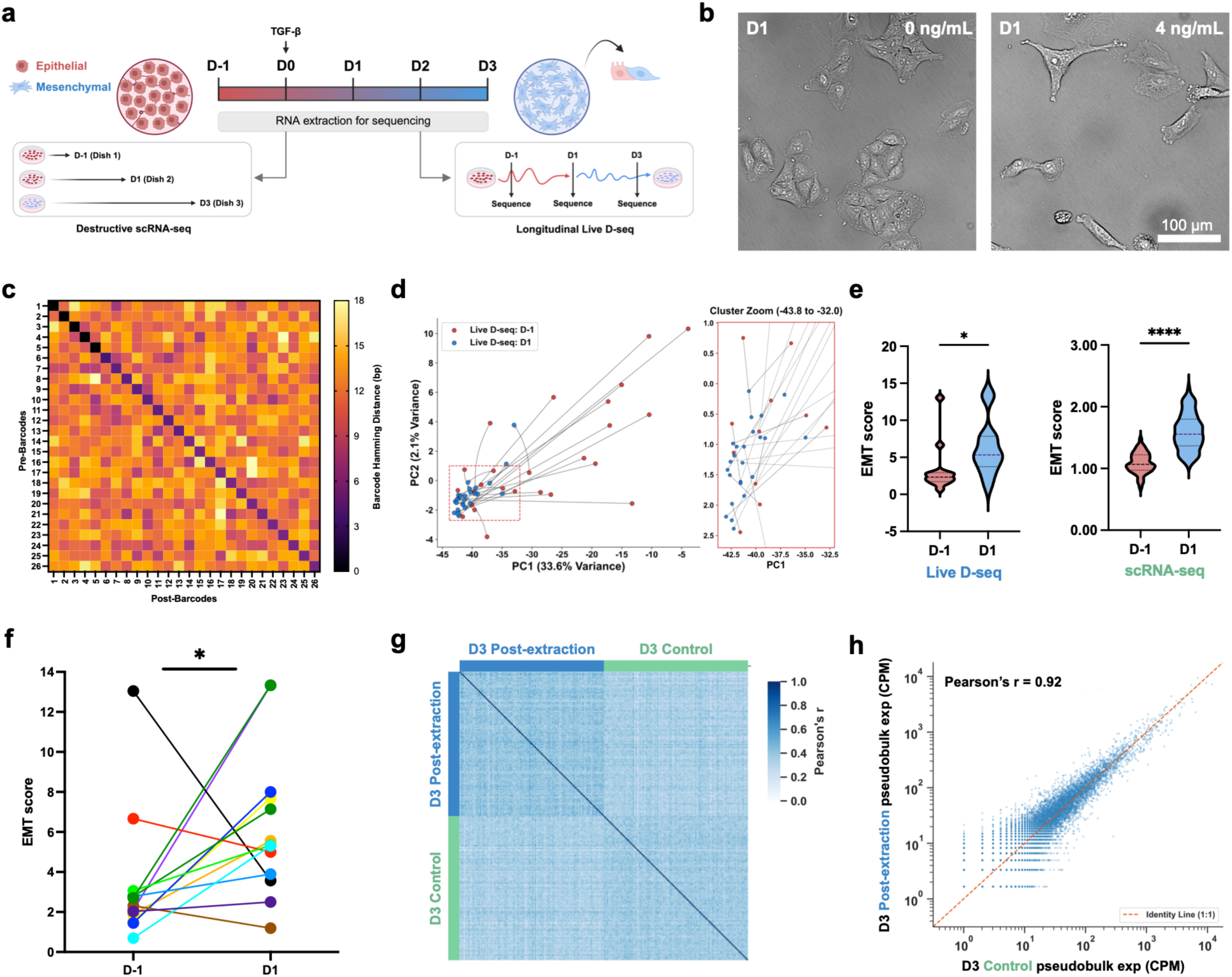
Live D-seq tracks EMT dynamics in single cells. (**a**) Experimental design. Barcoded A549 cells were treated with TGF-β at D0. Live D-seq sampled the same cell lineages at D-1 and D1, followed by full lysis at D3. scRNA-seq controls were lysed from separate wells at each time point. Created with BioRender.com. (**b**) Bright-field images of A549 cells at D1 without (0 ng/mL) or with (4 ng/mL) TGF-β. Scale bar: 100 µm. (**c**) Hamming distance between cellular barcodes recovered at D-1 (rows) and D1 (columns) for the 26 tracked cell lineages. (**d**) PCA of paired Live D-seq profiles at D-1 (red) and D1 (blue), with lines connecting the same cell lineage. The inset shows an enlarged view of the boxed region. (**e**) EMT scores at D-1 and D1 for Live D-seq (*n* = 13 tracked cell lineages) and scRNA-seq (*n* = 100 cells per time point), calculated from the 50 genes listed in **Supplementary Table 2**. Live D-seq cells with an EMT score of 0 at either time point, meaning no score genes were detected, were excluded. Statistical significance was assessed by a two-tailed ratio paired t-test (Live D-seq; *P* = 0.025) and a two-tailed lognormal Welch’s t-test (scRNA-seq; *P* < 0.0001). (**f**) EMT scores of individual tracked cell lineages from D-1 to D1 (*n* = 13 lineages, as in e). (**g**) Pairwise cell-cell Pearson correlation of A549 cells at D3 after two rounds of Live D-seq sampling compared with unsampled controls (100 highest-UMI cells per group). (**h**) Pseudobulk expression of previously sampled versus control cells at D3 (*n* = 132 previously sampled and 302 control cells).

Sequencing metrics and per-cell UMI and gene counts for both methods are shown in **Supplementary Fig. 7**. Principal component analysis (PCA) of the paired Live D-seq profiles showed a clear shift in transcriptional states between time points (**Fig. 4d**). The D-1 profiles were spread widely along PC1, while the D1 profiles clustered more closely together. Conventional scRNA-seq controls also showed separation between D-1 and D1 cells (**Extended Data Fig. 3a**). To quantify EMT, we first identified the 50 most strongly upregulated genes from D-1 to D1 in the scRNA-seq controls (**Supplementary Table 2**). These included canonical TGF-β and EMT-associated genes such as *SERPINE1*, *TGFBI*, *PMEPA1*, *COL4A1*, and *VIM*. We then used these 50 genes to calculate an EMT score for each cell in both datasets. EMT scores increased significantly from D-1 to D1 in both Live D-seq and scRNA-seq, demonstrating that non-destructively sampled RNA captures EMT-associated transcriptional changes consistent with those detected by scRNA-seq (**Fig. 4e**). By preserving cell viability, Live D-seq enabled longitudinal tracking of EMT responses within individual cell lineages, which is not possible with conventional scRNA-seq that requires cell lysis. EMT scores increased in most lineages, although the magnitude of the response varied (**Fig. 4f**). Baseline EMT scores also varied: a few cells already had high scores at D-1, before TGF-β treatment, and these cells showed little or no further increase, or a decrease, by D1. This fits the known heterogeneity of A549 cells, in which some cells show partial mesenchymal features even without stimulation.^38,39^

We next asked how closely Live D-seq agreed with scRNA-seq at each time point. Pseudobulk expression from the two methods was moderately correlated at both D-1 (Pearson’s *r* = 0.65) and D1 (Pearson’s *r* = 0.49; **Extended Data Fig. 3b,c**). Gene ontology enrichment also overlapped substantially at both time points (**Extended Data Fig. 3d,e**). Shared terms included responses to oxidative stress, regulation of cell migration and wound healing at D-1, and regulation of stress fiber assembly and actin monomer sequestration at D1, which are cytoskeletal processes associated with EMT. The most abundant genes also largely overlapped between methods at both time points, including *FTL*, *FTH1*, *ANXA2*, *TMSB4X*, *TMSB10*, *S100A6*, and *PPIA* (**Extended Data Fig. 3f,g**). At D1, TGF-β-responsive genes, including *TGFBI* in Live D-seq and *SERPINE1* in scRNA-seq, were among the most abundant transcripts, consistent with activation of TGF-β-associated transcriptional programs.

Finally, we asked whether repeated sampling disturbed the cells’ transcriptional state. At D3, we compared cells that had undergone two rounds of Live D-seq sampling with unsampled controls. The pairwise cell-cell correlation matrix showed little separation between the two groups. Between-group correlations (mean *r* = 0.34) were comparable to within-group correlations (mean *r* = 0.39 and 0.37 for repeatedly sampled and control cells, respectively; **Fig. 4g**). Pseudobulk expression was also highly correlated between repeatedly sampled and control cells (Pearson’s *r* = 0.92; **Fig. 4h**). Together, these results show that Live D-seq enables longitudinal tracking of transcriptional dynamics within the same cell lineages while preserving their overall transcriptomic profiles.

## 3. Discussion

Conventional scRNA-seq requires destructive cell lysis, while existing non-destructive single-cell sampling methods have limited scalability. To address both problems, we developed Live D-seq, a droplet-based platform that enables parallel RNA sampling from living cells for longitudinal single-cell transcriptomic profiling. Live D-seq integrates PTE, magnetic barcoded hydrogel beads, and hypotonic buffer-mediated RNA extraction to enable non-destructive transcriptomic profiling while preserving cell viability and maintaining transcriptional states. We demonstrated single-cell resolution and accurate cellular barcode assignment, with performance comparable to conventional scRNA-seq. By applying Live D-seq to TGF-β-induced EMT in A549 cells, we demonstrated longitudinal transcriptomic profiling of individual barcode-matched cell lineages, capturing increases in EMT scores consistent with conventional scRNA-seq despite sampling only a small fraction of intracellular RNA. Importantly, cells subjected to two rounds of RNA sampling maintained transcriptional profiles highly similar to unsampled controls, supporting the feasibility of repeated transcriptomic measurements in living cells.

Several limitations remain, although future improvements could further expand the capabilities of Live D-seq. Increasing barcode diversity through additional split-pool rounds or more barcode sequences per round could enable tracking of larger cell populations, particularly rare populations such as drug-resistant cells. Cell capture efficiency is currently limited by inefficient cell-hydrogel bead pairing during PTE, but the ability of PTE to rapidly generate droplets across a wide range of reaction volumes provides a route to processing more cells through larger reaction volumes or parallel reactions.^24^ This could compensate for losses that accumulate across sampling steps, increasing the number of cell lineages available for longitudinal tracking. In addition, because Live D-seq captures only a fraction of intracellular mRNA, it detects fewer genes per cell and preferentially captures abundant transcripts compared with conventional scRNA-seq. Improving library preparation could enhance transcriptomic sensitivity, particularly given the lower mapping efficiency observed with Live D-seq. Further optimization of RNA extraction conditions may also improve performance. Because extraction efficiency was measured in bulk without oil or vortexing, the effects of PTE on RNA recovery remain to be determined. Lastly, differences in membrane sensitivity across cell types, such as the limited benefit of saponin in A549 cells, highlight the need to optimize extraction conditions for individual cell types.

Despite these limitations, Live D-seq establishes a framework for parallel, non-destructive transcriptomic sampling with single-cell resolution that captures transcriptional changes consistent with conventional scRNA-seq while enabling longitudinal tracking of individual cell lineages. With improvements in capture efficiency and transcriptomic sensitivity, this approach could enable longitudinal analysis of larger cell populations and more sampling time points, providing new opportunities to investigate how early transcriptional states influence subsequent cell fate decisions. Such measurements could be particularly valuable for studying heterogeneous and transient responses during differentiation, drug resistance, and immune activation, where destructive snapshots cannot directly resolve transcriptional trajectories within individual cell lineages.

In summary, Live D-seq establishes a droplet-based platform for parallel, non-destructive RNA sampling and longitudinal tracking of individual cell lineages. By moving single-cell sequencing beyond destructive snapshots toward longitudinal measurements, this approach could facilitate studies of heterogeneous and transient cellular responses during differentiation, drug resistance, and immune activation.

## 4. Methods

### 4.1. Microfluidic chip fabrication

Microfluidic devices were designed in AutoCAD (Autodesk) and fabricated by conventional photolithography to produce a 70 µm-high SU-8 master mold. Polydimethylsiloxane (PDMS; Sylgard 184, Corning) was mixed at a 10:1 base-to-curing-agent ratio with 1.6% (w/w) iron powder (Thermo Fisher Scientific, 0073730) to suppress UV scattering within the channels during polymerization. The mixture was poured onto the master mold and cured overnight at 70 °C. Fluidic ports were punched with a 1 mm biopsy punch (Miltex). The bottom substrate was prepared by spin-coating PDMS onto glass slides and partially curing it at 60 °C for 12 min. The punched PDMS slab was placed onto the partially cured substrate, and the assembly was baked overnight at 60 °C for complete, leak-free bonding.

### 4.2. Magnetic hydrogel bead fabrication

Magnetic hydrogel beads were fabricated by droplet microfluidics with in-line photopolymerization.^25^ The hydrogel precursor (100 µL) contained 21.4% (v/v) poly(ethylene glycol) diacrylate (PEGDA; M_n_ 700, Sigma-Aldrich, 455008), 0.09% (w/v) lithium phenyl(2,4,6-trimethylbenzoyl)phosphinate (LAP; TCI, L0290), added from a 2.9% (w/v) stock, 10 µL of carboxyl magnetic particles (SPHERO Carboxyl Magnetic, High Iron, 0.24 µm, 1.0% w/v; Spherotech, CM-025-10H) and 50 µM acrydite-modified DNA oligonucleotide (sequence in **Supplementary Table 1**), made up to volume with TE buffer. For the cell-hydrogel bead pairing experiments, 1 mM acrylate-fluorescein (fluorescein O-acrylate; Sigma-Aldrich, 568856) was included in the precursor in place of the acrydite-modified DNA so that the hydrogel beads could be visualized by fluorescence microscopy.

Droplets were generated with syringe pumps (New Era Pump Systems), using QX200 Droplet Generation Oil for EvaGreen (Bio-Rad, 1864006) as the continuous phase. The precursor was driven by a syringe of 1× PBS as a spacer solution. To avoid oil backflow, flow was started at 1.0 mL/h (oil) and 0.8 mL/h (aqueous), then reduced to 0.6 mL/h and 0.1 mL/h, respectively, to produce droplets of ∼70 µm in diameter. Droplets were polymerized in the device outlet chamber by UV exposure through a 20× objective at 50% UV intensity on an IX83 inverted microscope (Olympus) with a DAPI filter set. We covered the droplet-formation junction with aluminum foil to prevent premature polymerization.

To recover the hydrogel beads, excess oil was removed after brief centrifugation. Then, 40 µL of 100% PFO (Thermo Fisher Scientific, B20156) and 200 µL of PBST (0.1% (v/v) Tween-20 in PBS) were added, and the mixture was pipetted for 30 s with low-retention tips. The aqueous phase was transferred to a new tube, diluted with 1 mL of PBST, and vortexed. After the hydrogel beads had settled, free magnetic particles were removed with the supernatant. Hydrogel beads were then centrifuged for 1 min at 13,500 × *g* and washed three times with PBST. Washed hydrogel beads were resuspended in 1 mL of PBST and stored at 4 °C until barcoding.

### 4.3. Split-pool barcoding of magnetic hydrogel beads

Hydrogel beads were barcoded by three rounds of split-pool splint ligation.^22,24^ For each round, barcode oligonucleotides and their complementary splint oligonucleotides (IDT; sequences in **Supplementary Table 1**) were reconstituted at 200 µM in IDTE buffer and mixed 1:1 (100 µM final). Each mix was annealed by heating to 72 °C for 2 min, then cooling to 22 °C at 0.1 °C/s, and stored at −20 °C. Third-round barcode oligonucleotides additionally contained a 12-nt UMI and an oligo(dT)_30_V capture sequence (V = A, C, or G).

All centrifugation steps were performed at 2,000 × *g* for 2 min unless otherwise stated. Before the first ligation, a splint oligonucleotide (PBB2) complementary to the hydrogel bead-bound acrydite oligonucleotide was annealed to the hydrogel beads. Hydrogel beads were washed once in TET buffer (10 mM Tris-HCl pH 8.0, 10 mM EDTA, 0.1% (v/v) Tween-20) and three times in pre-ligation (PL) buffer (10 mM Tris-HCl pH 7.5, 30 mM NaCl, 1 mM MgCl_2_, 0.1% (v/v) Tween-20). They were then incubated with 3.75 µM PBB2 in 1× T4 DNA ligase buffer, using a reaction volume twice the hydrogel bead pellet volume, annealed as above, and washed three times in TET.

For each ligation round, hydrogel beads were washed three times each in TET and PL and split equally into 40 tubes. Each tube received an equal volume of ligation mix containing 1× T4 DNA ligase buffer, 5 µM of one annealed barcode-splint duplex, and 20 U/µL T4 DNA ligase (NEB, M0202S). Ligation was performed at room temperature for 2 h, and the ligase was heat-inactivated at 65 °C for 10 min. Hydrogel beads were then pooled into TET and washed three times each in TET and PL.The PL wash was omitted after the final round. Three rounds of 40 barcodes yielded 40^3^ = 64,000 possible barcode combinations. The number of cells per run was limited to approximately one-fiftieth of the barcode diversity (∼1,280 cells) to keep the expected barcode collision rate at ∼1–2%. Accordingly, ∼1,200 cells were loaded per run.

After the third round, splint strands were removed to leave single-stranded barcodes. Hydrogel beads were washed once in nuclease-free water (3 min centrifugation), then incubated four times with 500 µL of denaturing solution (150 mM NaOH, 0.5% (w/v) Brij-35) for 3 min at room temperature with end-over-end rotation, with 3 min centrifugation between incubations. Hydrogel beads were then washed three times in low-salt (LS) buffer (10 mM Tris-HCl pH 8.0, 0.1 mM EDTA, 10 mM NaCl, 0.1% (v/v) Tween-20) and three times in TET, and stored at 4 °C.

To confirm barcoding, 10 µL of hydrogel beads were incubated with 1 µM FAM-labeled poly(A)_25_ oligonucleotide in 2× SSC (100 µL) for 20 min with shaking at 1,200 rpm. Hydrogel beads were then washed twice in PBST, centrifuged for 30 s at 3,380 × *g*, resuspended in 20–50 µL of PBST, and imaged on an IX83 inverted microscope. Images were analyzed in FIJI.

### 4.4. Cell culture

HEK293T (ATCC, CRL-3216), NIH-3T3 (ATCC, CRL-1658), and A549 cells (ATCC, CRM-CCL-185) were cultured in DMEM supplemented with 10% FBS and 1% penicillin-streptomycin. Jurkat cells (ATCC, TIB-152) were cultured in RPMI 1640 supplemented with 10% FBS and 1% penicillin–streptomycin. Primary human total T cells were provided by the Human Immunology Core and the Division of Transfusion Medicine and Therapeutic Pathology at the Perelman School of Medicine at the University of Pennsylvania. T cells were cultured in RPMI 1640 supplemented with 10% FBS, 1% penicillin–streptomycin, and 100 IU/mL recombinant human IL-2 (PeproTech, AF-200-02). All cells were maintained at 37 °C with 5% CO_2_.

### 4.5. Intracellular RNA extraction optimization

Adherent cells were detached with trypsin, and suspension cells were collected directly. Cells were washed in culture medium by centrifugation at 400 × g for 5 min. Aliquots of 50,000 cells were washed twice more in 1% BSA in DPBS without calcium and magnesium (0.22 µm filtered; 400 × g, 5 min) and resuspended in 50 µL of extraction buffer containing 0–100% (v/v) DI water in the same buffer. For saponin titrations, 0–100 µg/mL saponin was added to buffer containing 70% (v/v) DI water. After incubation for 30 min at room temperature, cells were pelleted at 500 × g for 5 min. Then 50 µL of supernatant was collected, purified, and buffer-exchanged into nuclease-free water using an RNA Clean & Concentrator-25 kit (Zymo Research, R1018), and eluted in 50 µL. For primary T cells, 40 µL was used in place of 50 µL for the extraction buffer, the collected supernatant, and the elution.

RNA concentration was measured with the Qubit RNA HS Assay Kit (Thermo Fisher Scientific, Q32855). Background signal was measured from extraction buffer processed identically without cells and subtracted from each sample. RNA extraction efficiency (%) was calculated as the background-corrected RNA mass (concentration × elution volume) divided by the estimated total RNA content of 50,000 cells. This estimate assumed ∼10 pg RNA per cell for cell lines^40^ and ∼2 pg per cell for primary T cells^41^. These are assumed reference values rather than measured per-cell RNA contents, so efficiencies are comparable across conditions but are not absolute, and are over-or underestimated where a cell type’s actual RNA content differs from the assumed value. Values are reported relative to the baseline condition: 0% (v/v) DI water for DI water titrations, and 0 µg/mL saponin for saponin titrations. Each condition was measured in three technical replicates.

### 4.6. Cell viability

Cell viability was assessed immediately after extraction (0 h) and after 24 h of culture, using a live/dead cell double staining kit (Sigma-Aldrich, 04511). For the 24 h time point, cells were returned to culture medium after extraction. Cells were incubated with calcein green AM (1 µM) and propidium iodide (1.5 µM) in DPBS containing calcium and magnesium for 15 min at 37 °C and imaged on an Echo Revolve (Echo) or IX83 inverted microscope. Live (calcein-positive) and dead (propidium iodide-positive) cells were counted in CellProfiler, FIJI, or manually for images with few cells. Viability was calculated for each replicate as live cells divided by total cells × 100, and averaged across three technical replicates.

### 4.7. Cell membrane staining

HEK293T cells were treated with extraction buffer (70% (v/v) DI water, 50 µg/mL saponin) and returned to culture for 24 h alongside untreated controls. Cells were washed three times with DPBS containing calcium and magnesium and stained with PE-conjugated anti-human CD9 antibody (BioLegend, 312105; clone HI9a; mouse IgG1κ) at 5 µg/mL for 25 min at 37 °C. After three washes with DPBS, nuclei were stained with Hoechst 33342 (Thermo Fisher Scientific, H3570) at a 1:10,000 dilution, followed by three further washes with DPBS. Images were acquired on an IX83 inverted microscope and analyzed in FIJI.

### 4.8. CloneSweeper cellular barcoding

Lentivirus was produced in HEK293T cells by polyethylenimine (PEI) transfection. PEI (PEI MAX, MW 40,000; Polysciences, 24765) was prepared at 1 mg/mL in water, adjusted to pH 7.0 with NaOH, and filter-sterilized through a 0.22 µm filter. HEK293T cells in 100 mm dishes at ∼75% confluence were transfected with the CloneSweeper transfer plasmid^21^ (a gift from the Shaffer laboratory), psPAX2, and pMD2.G at equal masses (3.33 µg each; 10 µg total plasmid per dish) using 50 µg PEI, corresponding to a PEI-to-plasmid mass ratio of 5:1. Plasmid and PEI were diluted separately in DPBS without calcium and magnesium, combined to a final volume of 1 mL, incubated for 10 min at room temperature, and added dropwise to 9 mL of culture medium. Medium was replaced the following day, and virus-containing supernatant was collected on days 3 and 5 post-transfection, filtered through a 0.22 µm filter, and stored at −80 °C.

A549 cells at ∼70% confluence were transduced at a 5:1 (v/v) medium-to-virus ratio in the presence of 0.8 µg/mL polybrene (Thermo Fisher Scientific, TR1003G). At 72 h post-transduction, eGFP-positive cells were isolated on a BD FACS Discover S8 cell sorter (BD Biosciences), with cells suspended in DPBS without calcium and magnesium, containing 1% BSA (0.22 µm filtered). Sorted cells were collected into culture medium and expanded for use in subsequent experiments.

### 4.9. Cellular barcode validation

For bulk barcode validation, RNA was extracted from barcoded and non-barcoded A549 cells with TRIzol reagent (Thermo Fisher Scientific, 15596018), using GlycoBlue coprecipitant (Thermo Fisher Scientific, AM9515) to visualize the pellet. cDNA was synthesized from RNA in a 20 µL reaction containing 1× Maxima RT buffer, 1 M betaine, 500 µM dNTPs, 1 U/µL RNase inhibitor, 5 µM oligo(dT)_30_, 1.25 µM biotinylated template-switching oligonucleotide, and 50 U of Maxima H Minus reverse transcriptase (Thermo Fisher Scientific, EP0751) on a SimpliAmp thermal cycler (Thermo Fisher Scientific) for 30 min at 50 °C and 5 min at 85 °C.

For on-hydrogel bead capture validation, barcoded magnetic hydrogel beads were incubated with total RNA from barcoded or non-barcoded A549 cells in 1× PBS for 30 min at room temperature. Hydrogel beads were washed three times in 2× RT buffer containing 1% (w/v) Pluronic F-127, and cDNA was synthesized on the hydrogel beads as described above for the lysed-cell controls. Each 10 µL qPCR reaction contained PowerUp SYBR Green Master Mix (Applied Biosystems, A25742), 1 ng of cDNA, and 500 nM each of forward and reverse primers (sequences in **Supplementary Table 1**). Reactions were run for 40 cycles on a QuantStudio 3 Real-Time PCR System (Applied Biosystems) using the manufacturer’s standard cycling conditions. Fold enrichment of the cellular barcode in barcoded samples relative to non-barcoded samples was calculated by the 2^−ΔΔCt^ method, with *ACTB* and *GAPDH* as controls and *RPL13A* for normalization. Each condition was measured in three technical replicates.

### 4.10. Single hydrogel bead and cell pairing optimization

Cells and hydrogel beads were co-encapsulated by particle-templated emulsification (PTE). Hydrogel beads were counted by microscopy: 5 µL of vortexed hydrogel bead suspension was diluted in 45 µL of PBST, and four 2 µL drops were counted on a coverslip. Hydrogel beads were washed three times in nuclease-free water containing 0.1% (w/v) Pluronic F-127 (3,380 × *g*, 30 s) and resuspended in 1% BSA in DPBS without calcium and magnesium.

Cells were detached with trypsin and stained with 1 µM calcein red-orange AM and Hoechst 33342 (1:10,000) in DPBS containing calcium and magnesium for 15 min at 37 °C, then washed and resuspended in 1% BSA in DPBS without calcium and magnesium at 250, 500, or 1,000 cells/µL. Excess buffer was removed from the hydrogel bead suspension to leave a pellet containing 10,000 hydrogel beads. Cells were added directly to the hydrogel bead pellet in a fixed volume of 2 µL and mixed by pipetting ten times, giving 500, 1,000, or 2,000 cells and cell-to-hydrogel bead ratios of 5%, 10%, and 20%. The mixture was overlaid with 500 µL of 2% (w/w) 008-FluoroSurfactant in HFE-7500 (RAN Biotechnologies, 008-FluoroSurfactant-2wtH-50G) and vortexed horizontally for 2 min (10 s at 3,200 rpm followed by 110 s at 2,500 rpm).

Droplets were imaged on an IX83 inverted microscope. Hydrogel beads were visualized by acrylate-fluorescein incorporated during fabrication. Five fields of view were imaged per condition, and droplets were scored in FIJI as containing one hydrogel bead with one cell, one hydrogel bead with more than one cell, or one cell without a hydrogel bead. Pairing efficiency was calculated for each field as the percentage of hydrogel beads paired with exactly one cell. A 10% cell-to-hydrogel bead ratio was used for all subsequent experiments.

### 4.11. Live D-seq pipeline

#### Droplet generation and RNA extraction

Barcoded hydrogel beads (12,000 beads) were washed three times in nuclease-free water containing 0.1% (w/v) Pluronic F-127 (3,380 × *g*, 30 s), washed once in hypotonic buffer, and resuspended in 50 µL of the same buffer containing 1 U/µL RNase inhibitor. Excess buffer was removed to leave a hydrogel bead pellet. Cells were detached with trypsin, washed twice in DPBS without calcium and magnesium containing 1% BSA (400 × g, 5 min), and resuspended at 600 cells/µL in the same buffer containing 1 U/µL RNase inhibitor. Then 2 µL of cell suspension (∼1,200 cells) was added to the hydrogel bead pellet and mixed by pipetting ten times, giving a final composition of 70% (v/v) DI water and 50 µg/mL saponin. Saponin was omitted for A549 cells. The mixture was overlaid with 500 µL of 2% (w/w) 008-FluoroSurfactant in HFE-7500 and vortexed horizontally for 2 min (10 s at 3,200 rpm, then 110 s at 2,500 rpm). Droplets were incubated for 25 min at room temperature.

#### Demulsification and cell recovery

Excess oil was removed, and the droplet phase was transferred to a PCR tube. Droplets were broken by electro-coalescence with a Zerostat anti-static gun (VWR, Z108812), and the oil phase was removed. Hydrogel beads and cells were eluted in 50 µL of serum-free DMEM containing 0.1% (w/v) Pluronic F-127 and transferred to a PCR tube on a magnetic rack. After 30–60 s, 90–95% of the supernatant, containing the cells, was transferred to a 384-well plate and diluted with the same volume of medium containing 20% FBS. Cells were placed immediately at 37 °C, and the medium was replaced the following day. Hydrogel beads were retained for library preparation.

#### scRNA-seq controls

For scRNA-seq controls, cells were resuspended in DPBS without calcium and magnesium, containing 0.04% BSA and combined with barcoded hydrogel beads in lysis buffer (29 U/mL proteinase K, 70 mM DTT), giving a final composition of ∼19–21 U/mL proteinase K and ∼47–50 mM DTT after dilution by the cell suspension. Droplets were generated as above, and cells were lysed in the emulsion at 65 °C for 35 min with the lid at 105 °C. Excess oil was removed, 40 µL of 100% PFO was layered on the emulsion, followed by 200 µL of high-salt buffer (250 mM Tris-HCl pH 8.0, 375 mM KCl, 15 mM MgCl_2_, 50 mM DTT). The mixture was pipetted for 30 s with low-retention tips. The aqueous phase was transferred to a new 1.5 mL tube.

#### Reverse transcription

Hydrogel beads were washed three times in 300 µL of 2× RT buffer (100 mM Tris-HCl pH 8.3, 150 mM KCl, 6 mM MgCl_2_, 20 mM DTT) containing 1% (w/v) Pluronic F-127. Reverse transcription was performed in a 50 µL reaction containing 1× Maxima RT buffer, 1 M betaine, 200 µM dNTPs, 4% (w/v) Ficoll PM-400, 1 U/µL RNase inhibitor, 1.25 µM biotinylated template-switching oligonucleotide, and 0.5 U/µL Maxima H Minus reverse transcriptase. 1 U/µL reverse transcriptase was used for the lysed-cell controls. Reactions were incubated in a thermal cycler for 30 min at room temperature, then 90 min at 42 °C, with vortexing every 30 min. Hydrogel beads were then washed once each in TE containing 0.5% (w/v) SDS, TE containing 0.01% (v/v) Tween-20, and TE containing 0.05% (w/v) Pluronic F-127.

#### cDNA amplification and library preparation

cDNA was amplified in a 50 µL reaction containing 1× KAPA HiFi HotStart ReadyMix (Roche, KK2602) and 0.25 µM template-switch PCR primer, with cycling at 95 °C for 3 min, then 23 cycles (Live D-seq) or 12 cycles (scRNA-seq controls) of 98 °C for 15 s, 67 °C for 20 s and 68 °C for 4 min, followed by 72 °C for 5 min. Amplified cDNA was purified with two rounds of SPRI selection (0.8× then 0.6×) and eluted in 12 µL of nuclease-free water. cDNA size distribution was assessed on a TapeStation D5000 assay (Agilent), and concentration was measured by Qubit.

Transcriptome libraries were prepared from 1 ng of cDNA with the Nextera XT DNA Library Preparation Kit (Illumina, FC-131-1024) using 0.5 µM each of custom P5 and P7 primers, purified with 0.7× SPRI beads and eluted in 15 µL. Cellular barcode libraries were prepared without tagmentation in a 50 µL reaction containing 1× KAPA HiFi HotStart ReadyMix, 1 ng of cDNA, 0.5 µM P7 primer and 0.25 µM P5 primer, with cycling at 95 °C for 3 min; 10 cycles of 95 °C for 20 s, 70 °C for 30 s (−0.2 °C per cycle) and 72 °C for 20 s; 8 cycles of 95 °C for 20 s, 68 °C for 30 s and 72 °C for 20 s; and 72 °C for 4 min. Libraries were purified by double-sided SPRI selection (0.5×/0.8×) and eluted in 15 µL.

### 4.12. Transcriptome perturbation assessment

HEK293T cells were processed by Live D-seq as described above, using non-barcoded hydrogel beads. Hydrogel beads were discarded, and recovered cells were returned to culture for 48 h. Treated cells and untreated controls were then profiled using the droplet-based scRNA-seq protocol described above, with barcoded hydrogel beads and in-droplet lysis. The two conditions were processed side by side as separate emulsions, with ∼1,200 cells loaded per condition, and sequenced as separate libraries.

### 4.13. Cross-species experiment

HEK293T (human) and NIH-3T3 (mouse) cells were mixed at a 1:1 ratio and processed by Live D-seq as described above, with barcoded hydrogel beads and ∼1,200 cells loaded. A parallel sample of the same cell mixture was processed by droplet-based scRNA-seq with in-droplet lysis as a reference. Hydrogel beads from both samples were sequenced as separate libraries. Species assignment and doublet calling are described below.

### 4.14. Cellular barcode and hydrogel bead barcode pairing

Barcoded A549 cells were processed by Live D-seq as described above, with barcoded hydrogel beads and ∼1,200 cells loaded. Saponin was omitted from the extraction buffer, as it provided negligible improvement in RNA extraction from A549 cells. A parallel sample was processed by droplet-based scRNA-seq with in-droplet lysis as a reference. Both transcriptome and cellular barcode libraries were prepared as described above and sequenced as separate libraries. Assignment of cellular barcodes to hydrogel beads and calculation of barcode purity are described below.

### 4.15. EMT time course

Sorted barcoded A549 cells were seeded in 48-well plates at ∼70% confluence and cultured for 48 h. At D-1, cells were detached from a single well and sampled by Live D-seq as described above, with saponin omitted: 4,800 cells were processed across four parallel reactions (∼1,200 cells each). Recovered cells were returned to culture in a 384-well plate, and TGF-β (R&D Systems, 7754-BH) was added at D0 to a final concentration of 4 ng/mL. At D1, cells were sampled again across two parallel reactions (2,400 cells total) and returned to culture. At D3, the remaining cells were fully lysed and processed by droplet-based scRNA-seq with in-droplet lysis. Unsampled control cells were cultured in parallel and lysed at D3 alongside the previously sampled cells. As additional controls, separate wells of barcoded A549 cells were lysed and processed by droplet-based scRNA-seq at each time point.

Cells were imaged at each time point on an IX83 inverted microscope to monitor eGFP expression and morphology. To confirm the TGF-β dose, A549 cells were treated with 0 or 4 ng/mL TGF-β and imaged by bright-field microscopy at D1 on the same instrument. Barcode matching across time points and EMT scoring are described below.

### 4.16. Sequencing and read processing

Libraries were sequenced on an AVITI system (Element Biosciences) with paired-end reads (Read 1, 99 nt; Read 2, 199 nt) using the AVITI 2 × 150 Sequencing Kit Cloudbreak FS. Reads were demultiplexed by sample index with bases2fastq (v2.3.0). In all libraries, Read 1 contains the hydrogel bead barcode and the unique molecular identifier (UMI). Read 2 contains the cDNA in transcriptome libraries and the cellular barcode in co-barcoding libraries.

Hydrogel bead barcodes and UMIs were extracted from Read 1 with custom Python scripts. Each hydrogel bead barcode consists of three 8-nt split-pool segments separated by fixed linkers: GATCT, a 0–3-nt spacer, segment 1, ATGCATC, segment 2, CCTCGAG, segment 3, then a 12-nt UMI. Each linker was located at its expected position, allowing one mismatch, and reads missing any linker were discarded. The three segments were concatenated into a 24-nt hydrogel bead barcode and written, with the UMI, to a synthetic barcode read. In transcriptome libraries, Read 2 was trimmed at the first poly(A) stretch (8 A), TruSeq or Nextera adapter sequence, or hydrogel barcode oligonucleotide sequence (identified by its linker–barcode–linker junctions in either orientation). Reads shorter than 20 nt after trimming were discarded.

Trimmed reads were aligned and counted with STARsolo (STAR v2.7.11b) using a 24-nt cell barcode, a 12-nt UMI and a whitelist of all 64,000 valid hydrogel bead barcodes (40 × 40 × 40 combinations). Alignment was permissive (--outFilterScoreMinOverLread 0.1, -- outFilterMatchNminOverLread 0.45). Only uniquely mapped reads were counted. Reads were assigned to genes including introns (GeneFull), with STARsolo’s default barcode correction (one mismatch to the whitelist) and UMI collapsing. Human samples were aligned to GRCh38 (primary assembly) with the GENCODE v45 annotation. Human–mouse (cross-species) samples were aligned to a combined GRCh38 (GENCODE v45) and GRCm39 (GENCODE vM34) reference. Sequencing saturation, read number, and the fraction of uniquely mapped reads were taken from the STARsolo summary statistics.

Cells were called in three ways, depending on the experiment. In the cross-species experiment, hydrogel beads with at least 500 UMIs (scRNA-seq) or 24 UMIs (Live D-seq) in the raw STARsolo matrix were classified. For cell–cell and pseudobulk comparisons and for scRNA-seq controls, cells were called with STARsolo’s EmptyDrops_CR filter (default parameters), and only cells with at least 500 UMIs were used for pseudobulk and differential expression. For the EMT time course, hydrogel beads with at least 10 UMIs in the raw STARsolo matrix were kept for pairing with cellular barcodes, as described below.

### 4.17. Single-cell analysis

Unless stated otherwise, log denotes the natural logarithm.

#### Species assignment

For each hydrogel bead above the UMI cutoff, UMIs were summed separately over human (ENSG) and mouse (ENSMUSG) genes. Hydrogel beads with at least 85% of UMIs from one species were assigned to that species; all others were called doublets. All downstream cross-species analyses used each species’ singlets and counted only that species’ own genes.

#### Cellular barcode extraction and purity

In co-barcoding libraries, the 20-nt cellular barcode was extracted from Read 2 between a 23-nt fixed prefix and a 7-nt anchor, each allowing one mismatch at its expected position. The hydrogel bead barcode was extracted from Read 1 as above. Sequencing errors were corrected in two steps: first, across the dataset, cellular barcodes within one mismatch of a barcode with at least 5-fold more reads were merged into it; second, within each cellular barcode, hydrogel bead barcodes within two mismatches of a hydrogel bead with at least 2-fold more reads (and at least 5 reads) were merged. Barcode purity was the fraction of a hydrogel bead’s co-barcoding read pairs assigned to its most abundant (dominant) cellular barcode. In the cellular barcode and hydrogel bead barcode pairing experiment, hydrogel beads whose dominant barcode had at least 100 (scRNA-seq) or 15 (Live D-seq) read pairs were analyzed, and hydrogel beads with at least 85% purity were called singlets. In the EMT time course, a hydrogel bead was paired with a cellular barcode only if its dominant barcode had at least one read pair and at least 70% purity, and the pairing was reciprocal (the hydrogel bead was also the best-supported hydrogel bead for that cellular barcode).

Transcriptome hydrogel beads were matched to co-barcoding hydrogel beads by exact barcode match, otherwise by the nearest hydrogel bead within two mismatches (Hamming distance). Ties were left unassigned.

#### Lineage matching across time points

Each D1 cellular barcode was matched to a D-1 barcode by exact match, otherwise to the nearest D-1 barcode within five mismatches (Hamming distance over the 20-nt barcode). D1 barcodes with several equally close D-1 barcodes were left unmatched, and when several D1 barcodes matched one D-1 barcode, the nearest was kept. A lineage was removed if its own D-1–D1 pairing fit worse (by Hamming distance) than a cross-pairing with another lineage, and lineages without transcriptome data at either time point were removed. The number of matches expected by chance at each distance was estimated by matching D1 barcodes to shuffled barcodes (200 shuffles).

#### Transcript composition and gene detection

Transcript composition is the share of pseudobulk counts per million (CPM) in each GENCODE gene category: mitochondrial genes (chrM); ribosomal protein genes (protein-coding RPL/RPS genes); lncRNA; other protein-coding genes; and other genes. The mitochondrial fraction per cell used the same chrM gene set. Pseudobulk profiles summed raw UMI counts over all singlets of a species. For gene detection, a gene was detected in an assay if it had at least one UMI in that assay’s pseudobulk. For each group of genes (detected only in scRNA-seq, or in both assays), the average scRNA-seq UMI count per gene was calculated.

#### Most abundant genes

For each assay (and time point, where applicable), the most abundant genes by pseudobulk CPM were selected: 10 per assay in the cross-species experiment and 15 per assay in the EMT time course. These were grouped by whether they were among the most abundant in Live D-seq only, scRNA-seq only, or both. The list was topped up with the next most abundant genes detected in both assays, ranked by mean log(CPM + 1), to twice the per-assay number. Genes tied at the cutoff were all kept. Unannotated genes (Ensembl IDs, LOC and LINC, plus mouse Gm genes in the cross-species experiment) and mitochondrial genes were excluded, and ribosomal protein genes were excluded where stated. Expression is shown as natural log(CPM + 1).

#### Pseudobulk correlations

Pseudobulk expression was calculated as CPM of raw UMI counts summed over all cells in each group. Pearson’s *r* was calculated on log_10_ CPM over genes detected in both groups (CPM > 0 in both), so genes absent from either group were excluded. The comparisons were: post-extraction versus control cells 48 h after sampling in the transcriptome perturbation assessment; previously sampled versus control cells at D3 in the EMT time course; Live D-seq versus scRNA-seq singlets in the cross-species experiment, per species; and the 26 tracked Live D-seq lineages versus the 100 highest-UMI scRNA-seq cells at each time point in the EMT time course.

#### Cell-cell correlation heatmaps

The highest-UMI cells per group were selected: 150 per group in the transcriptome perturbation assessment and 100 per group at D3 in the EMT time course. Each cell was randomly downsampled (binomial, fixed seed) to the UMI total of the shallowest selected cell, so that all cells were compared at equal depth. Expression was normalized to log(counts per 10,000 + 1). Ribosomal and mitochondrial genes were excluded. Pearson correlations between all pairs of cells were calculated over the genes detected in at least 50% of the pooled cells.

#### PCA

The Live D-seq profiles of the 26 tracked lineages at D-1 and D1 were combined with the 100 highest-UMI scRNA-seq cells at each time point. Expression was normalized to log(counts per 10,000 + 1). The 1,000 genes with the highest variance across all cells were scaled to zero mean and unit variance per gene. A single principal component analysis (scikit-learn) was fitted on all cells together, and the Live D-seq and scRNA-seq cells are plotted separately from this shared analysis.

#### EMT score

The EMT gene set (**Supplementary Table 2**) is the 50 genes with the highest log_2_ fold change between scRNA-seq cells at D1 and D-1 (cells with at least 500 UMIs), excluding unannotated and mitochondrial genes. The fold change was the difference in mean log(counts per 10,000 + 1) between time points, divided by ln 2. All 50 genes were significant (two-tailed Welch’s t-test per gene, Benjamini-Hochberg adjusted *P* < 0.05). Each cell’s EMT score was the percentage of its detected genes that belong to the EMT gene set. A score of 0 means no gene from the set was detected.

#### Gene Ontology enrichment

GO enrichment was performed with Enrichr (GO Biological Process 2021 library) through GSEApy (v1.3.1). For each assay and time point, the 50 most abundant genes in the pseudobulk were tested against Enrichr’s default background. Unannotated, ribosomal protein, and mitochondrial genes were excluded, and genes tied at the cutoff were left out. Enrichment was tested with Fisher’s exact test, with Benjamini–Hochberg correction; terms with adjusted *P* < 0.05 were considered significant. For each assay, the 10 terms with the lowest adjusted *P* were shown, topped up to 20 terms per time point.

### 4.18. Statistics

Unless stated otherwise, *n* is the number of individual cells (hydrogel beads) and technical replicates are stated in the figure legends. Live D-seq EMT scores were compared between D-1 and D1 with a two-tailed ratio paired t-test, paired by tracked lineage. The ratio test uses log-transformed scores, so 13 of the 26 tracked lineages that had an EMT score of 0 at one or both time points were excluded (*n* = 13). scRNA-seq EMT scores were compared between time points with a two-tailed Welch’s t-test on log-transformed scores (lognormal Welch’s t-test), with *n* = 100 cells per time point.

These tests were performed in GraphPad Prism (v11.1.0, GraphPad Software); all other analyses were performed in Python. Significance was set at *P* < 0.05. For differential expression and GO enrichment, *P* values were adjusted for multiple testing with the Benjamini–Hochberg method. Pearson correlations are reported as descriptive measures of similarity, without *P* values. Data from the pairing, recovery, extraction, and viability optimizations are shown as mean ± s.d. of three replicates.

## 6. Acknowledgements

The data for this manuscript were generated in the Penn Cytomics and Cell Sorting Shared Resource Laboratory at the University of Pennsylvania (RRID: SCR_022376). Penn Cytomics is partially supported by the Abramson Cancer Center NCI Grant (P30 CA016520). We thank the Singh Center for Nanotechnology for fabrication support of the microfluidic devices. We also thank Emily Cento, Zhilin Chen, Max A. Eldabbas, and Emileigh Maddox of the Human Immunology Core (HIC) and the Division of Transfusion Medicine and Therapeutic Pathology at the Perelman School of Medicine at the University of Pennsylvania for providing de-identified primary human total T cells that were purified from healthy donor apheresis using RosetteSep kits (STEMCELL Technologies). The HIC is supported in part by NIH P30 AI045008 and P30 CA016520. HIC RRID: SCR_022380.

## 8. Author information

These authors contributed equally: David E. Reynolds, Sangmo Lee, and Daniel Oh

## 9. Ethics declarations

Competing interests

J.K. is a scientific cofounder and adviser for Aperture Bio and Cellfoundry Bio. The remaining authors declare no competing interests.

---

**Extended Data Figure 1:**
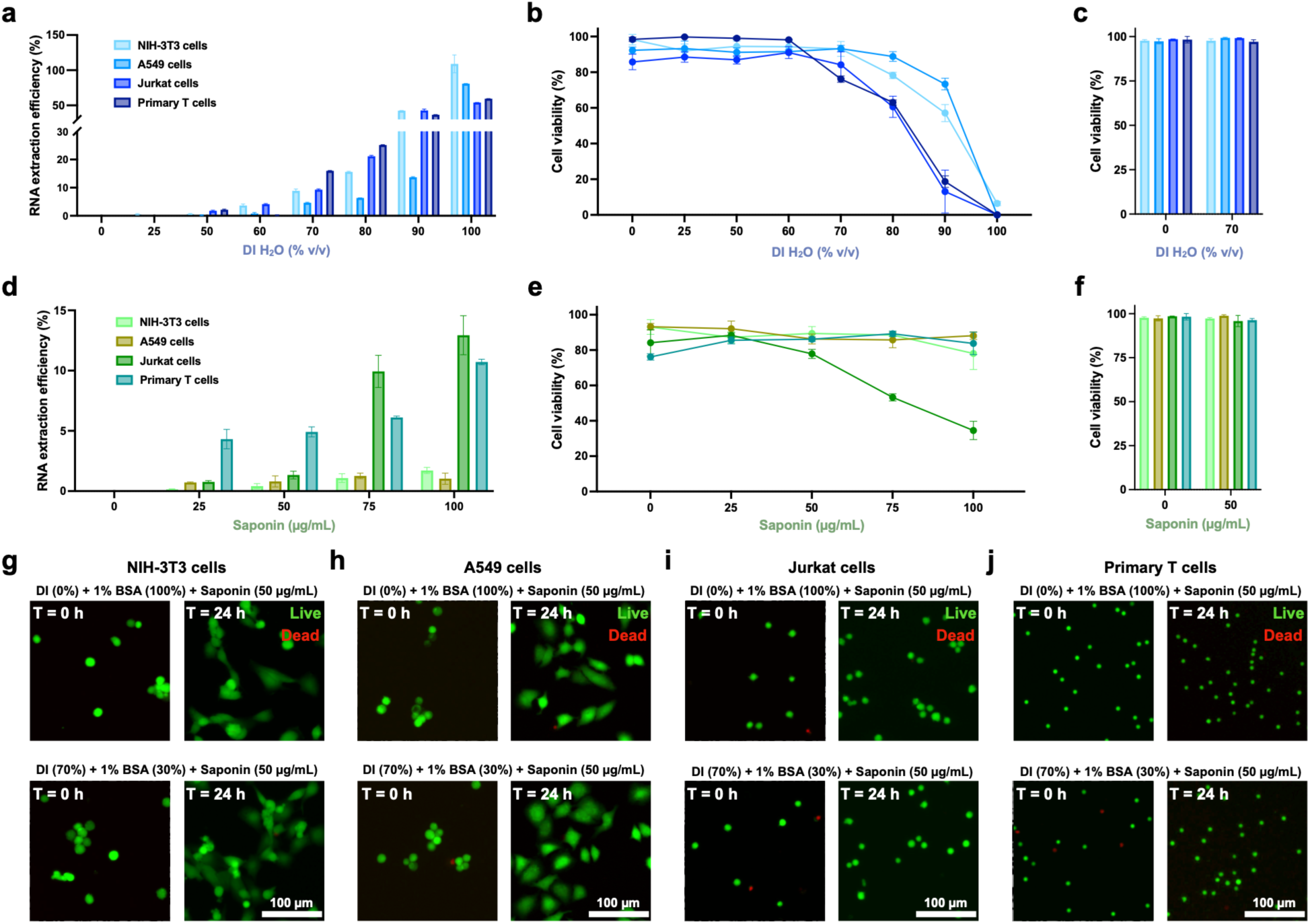
Live-cell subsampling of different cell types. (**a–c**) RNA extraction efficiency (**a**), immediate cell viability (**b**), and viability at 24 h (**c**) of NIH-3T3, A549, Jurkat, and primary T cells across deionized (DI) water concentrations, without saponin. Extraction efficiency in (**a**) is shown relative to 0% (v/v) DI water. (**d–f**) Additional RNA extraction efficiency (**d**), immediate cell viability (**e**), and viability at 24 h (**f**) across saponin concentrations in 70% (v/v) DI water. Extraction efficiency in (**d**) is shown relative to 0 µg/mL saponin. (**g–j**) Live/dead staining of NIH-3T3 (**g**), A549 (**h**), Jurkat (**i**), and primary T cells (**j**) at 0 and 24 h after treatment with control buffer (0% (v/v) DI water) or extraction buffer (70% (v/v) DI water), both containing 1% bovine serum albumin (BSA) and 50 µg/mL saponin. Live cells were stained with calcein green AM (green) and dead cells with propidium iodide (red). Data in (**a–f**) are shown as mean ± s.d. (*n* = 3 technical replicates). Scale bars: 100 µm.

**Extended Data Figure 2:**
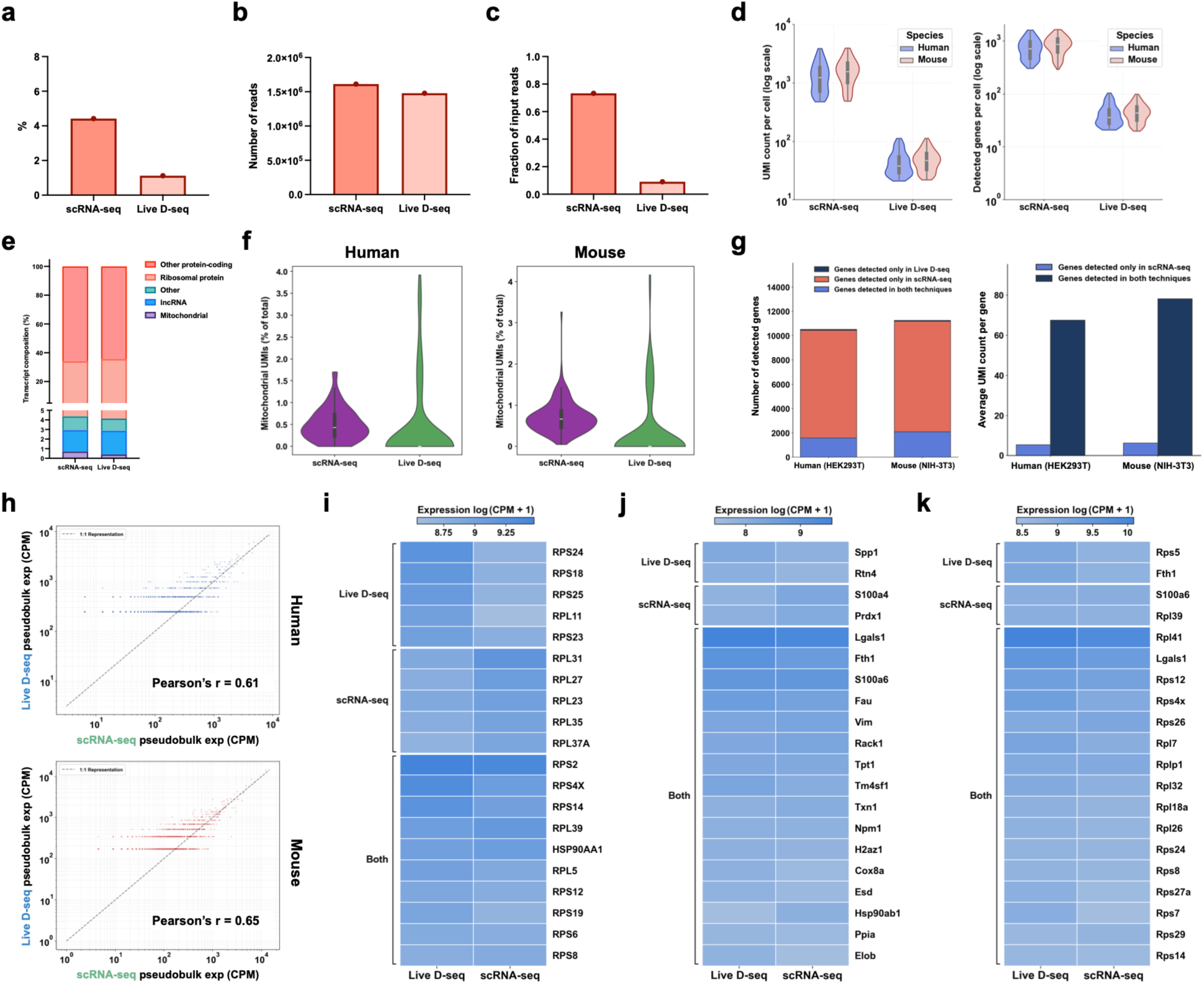
Cross-species additional analysis. (**a–c**) Sequencing saturation (**a**), number of sequencing reads (**b**), and fraction of input reads uniquely mapped (**c**) for single-cell RNA sequencing (scRNA-seq) and Live D-seq libraries. Each point represents one library. (**d–k**) Analyses include only hydrogel beads above 500 UMIs (scRNA-seq) or 24 UMIs (Live D-seq), as in Fig. 3a. (**d**) UMI counts (left) and detected genes (right) per cell for human and mouse cells, shown on a log_10_ scale. (**e**) Transcript composition of mouse cells by gene category. (**f**) Fraction of mitochondrial UMIs per cell in human (left) and mouse (right) cells. (**g**) Number of genes detected by Live D-seq only, scRNA-seq only, or both (left), and average UMI count per gene in scRNA-seq for genes detected only in scRNA-seq or in both methods (right). In (**d–g**), UMIs and genes were counted across all of that species’ genes, without gene filtering. (**h**) Pseudobulk expression from Live D-seq versus scRNA-seq for human (top) and mouse (bottom) cells. (**i–k**) Expression of the most abundant genes, grouped by whether they were among the most abundant in Live D-seq only, scRNA-seq only, or both, for human cells, including ribosomal protein genes (**i**), and mouse cells, excluding (**j**) or including (**k**) ribosomal protein genes.

**Extended Data Figure 3:**
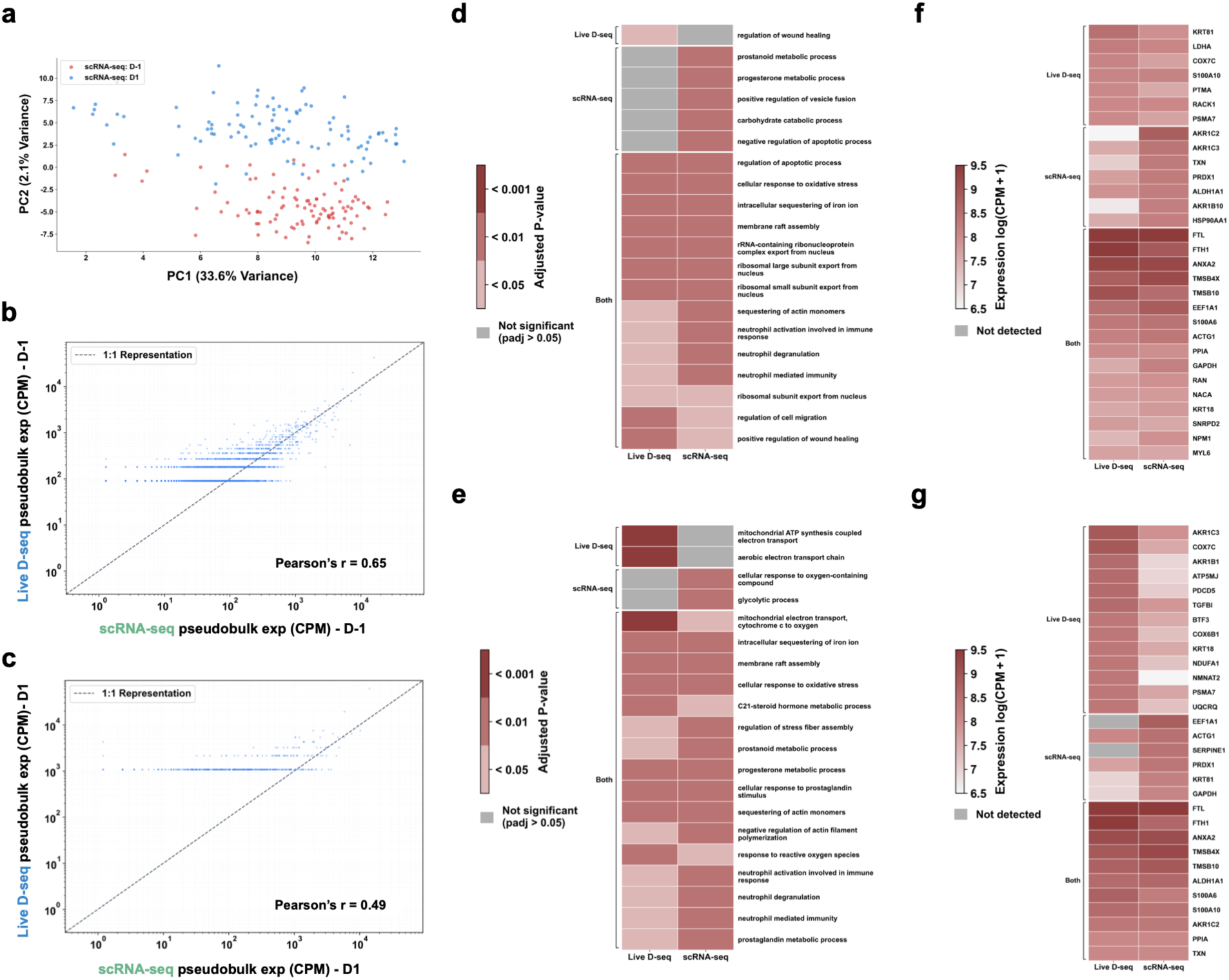
Additional analysis of EMT dynamics tracked by Live D-seq. (**a**) scRNA-seq control cells at D-1 (red) and D1 (blue), from the combined PCA shown in Fig. 4d. (**b,c**) Pseudobulk expression from Live D-seq versus scRNA-seq at D-1 (**b**) and D1 (**c**). (**d,e**) Gene ontology (GO) biological process terms enriched among the 50 most abundant genes in Live D-seq only, scRNA-seq only, or both, at D-1 (**d**) and D1 (**e**), colored by Benjamini-Hochberg adjusted *P* value; gray indicates terms that were not significant (adjusted *P* > 0.05). (**f,g**) Expression of the most abundant genes, grouped by whether they were among the most abundant in Live D-seq only, scRNA-seq only, or both, at D-1 (**f**) and D1 (**g**). Gray indicates genes that were not detected. Ribosomal protein genes were excluded from (**d–g**). Analyses include the 26 lineage-tracked Live D-seq cells and 100 scRNA-seq cells per time point.

## References

1. Papalexi, E. & Satija, R. Single-cell RNA sequencing to explore immune cell heterogeneity. Nat Rev Immunol 18, 35–45 (2018).

2. Stuart, T. & Satija, R. Integrative single-cell analysis. Nat Rev Genet 20, 257–272 (2019).

3. Li, X. & Wang, C.-Y. From bulk, single-cell to spatial RNA sequencing. Int J Oral Sci 13, 36 (2021).

4. Jolasun, Y., Song, K., Zheng, Y., Wang, J., Fonseca, G. J., Eidelman, D. H. & Ding, J. SIDISH integrates single-cell and bulk transcriptomics to identify high-risk cells and guide precision therapeutics through in silico perturbation. Nat Commun 16, 11271 (2025).

5. Van de Sande, B., Lee, J. S., Mutasa-Gottgens, E., Naughton, B., Bacon, W., Manning, J., Wang, Y., Pollard, J., Mendez, M., Hill, J., Kumar, N., Cao, X., Chen, X., Khaladkar, M., Wen, J., Leach, A. & Ferran, E. Applications of single-cell RNA sequencing in drug discovery and development. Nat Rev Drug Discov 22, 496–520 (2023).

6. Cuomo, A. S. E., Seaton, D. D., McCarthy, D. J., Martinez, I., Bonder, M. J., Garcia-Bernardo, J., Amatya, S., Madrigal, P., Isaacson, A., Buettner, F., Knights, A., Natarajan, K. N., HipSci Consortium, Vallier, L., Marioni, J. C., Chhatriwala, M. & Stegle, O. Single-cell RNA-sequencing of differentiating iPS cells reveals dynamic genetic effects on gene expression. Nat Commun 11, 810 (2020).

7. Zheng, H., Maestri, E., Moore, A. R., Ganesan, A., Enriquez, J., Durán, V., Einav, S., van der Poll, T., Bouma, H. R., Atreya, M. R., Efron, P. A., Moldawer, L. L., Sevransky, J., Wright, D. W., Martin-Loeches, I., Karvunidis, T., Hinson, J., Hager, D. N., Rothman, R. E., Giamarellos-Bourboulis, E. J., Sweeney, T. E., Rogers, A. J., Gombar, S., Maddali, M. V., Chen, J. H., Shah, N. & Khatri, P. A single-cell atlas of human neutrophils reveals neutrophil heterogeneity across health and disease. Nat Immunol 27, 2322–2334 (2026).

8. Jovic, D., Liang, X., Zeng, H., Lin, L., Xu, F. & Luo, Y. Single-cell RNA sequencing technologies and applications: a brief overview. Clin Transl Med 12, e694 (2022).

9. Reynolds, D. E., Roh, Y. H., Oh, D., Vallapureddy, P., Fan, R. & Ko, J. Temporal and spatial omics technologies for 4D profiling. Nat Methods 22, 1408–1419 (2025).

10. Reynolds, D. E., Roh, Y. H., Chintapula, U., Huynh, E., Vallapureddy, P., Tran, H. H., Lee, D., Allen, M. G., Xu, X. & Ko, J. Vertically Aligned Nanowires for Longitudinal Intracellular Sampling. ACS Nano 19, 13073–13083 (2025).

11. Chen, W., Guillaume-Gentil, O., Rainer, P. Y., Gäbelein, C. G., Saelens, W., Gardeux, V., Klaeger, A., Dainese, R., Zachara, M., Zambelli, T., Vorholt, J. A. & Deplancke, B. Live-seq enables temporal transcriptomic recording of single cells. Nature 608, 733–740 (2022).

12. Guillaume-Gentil, O., Grindberg, R. V., Kooger, R., Dorwling-Carter, L., Martinez, V., Ossola, D., Pilhofer, M., Zambelli, T. & Vorholt, J. A. Tunable Single-Cell Extraction for Molecular Analyses. Cell 166, 506–516 (2016).

13. Marcuccio, F., Chau, C. C., Tanner, G., Elpidorou, M., Finetti, M. A., Ajaib, S., Taylor, M., Lascelles, C., Carr, I., Macaulay, I., Stead, L. F. & Actis, P. Single-cell nanobiopsy enables multigenerational longitudinal transcriptomics of cancer cells. Sci Adv 10, eadl0515 (2024).

14. Cai, L., Lin, S., Qiu, M., Lin, L., Li, F., Liu, J., Zou, Y., Na, X., Liang, S., Xu, X., Yang, C. & Li, J. scBiopsy-seq: a platform for temporal single-cell RNA-seq analysis. bioRxiv 2025.03.26.645409 (2025).

15. Lovatt, D., Ruble, B. K., Lee, J., Dueck, H., Kim, T. K., Fisher, S., Francis, C., Spaethling, J. M., Wolf, J. A., Grady, M. S., Ulyanova, A. V., Yeldell, S. B., Griepenburg, J. C., Buckley, P. T., Kim, J., Sul, J.-Y., Dmochowski, I. J. & Eberwine, J. Transcriptome in vivo analysis (TIVA) of spatially defined single cells in live tissue. Nat Methods 11, 190–196 (2014).

16. Quillin, A. L., Karloff, D. B., Ayele, T. M., Flores, T. F., Chen, G., McEachin, Z. T., Valdez-Sinon, A. N. & Heemstra, J. M. Imaging and Tracking RNA in Live Mammalian Cells via Fluorogenic Photoaffinity Labeling. ACS Chem Biol 20, 707–720 (2025).

17. Pfeiffer, P., Nilsson, J. R., Gallud, A., Baladi, T., Le, H.-N., Bood, M., Lemurell, M., Dahlén, A., Grøtli, M., Esbjörner, E. K. & Wilhelmsson, L. M. Metabolic RNA labeling in non-engineered cells following spontaneous uptake of fluorescent nucleoside phosphate analogues. Nucleic Acids Res 52, 10102–10118 (2024).

18. Wang, D., Shalamberidze, A., Arguello, A. E., Purse, B. W. & Kleiner, R. E. Live-Cell RNA Imaging with Metabolically Incorporated Fluorescent Nucleosides. J Am Chem Soc 144, 14647–14656 (2022).

19. Wang, H., Nakamura, M., Abbott, T. R., Zhao, D., Luo, K., Yu, C., Nguyen, C. M., Lo, A., Daley, T. P., La Russa, M., Liu, Y. & Qi, L. S. CRISPR-mediated live imaging of genome editing and transcription. Science 365, 1301–1305 (2019).

20. Xia, C., Colognori, D., Jiang, X. S., Xu, K. & Doudna, J. A. Single-molecule live-cell RNA imaging with CRISPR-Csm. Nat Biotechnol 43, 2023–2030 (2025).

21. Vander Velde, R. J., Ng, R. W. S., Coté, C. & Shaffer, S. M. Multiplexed enrichment and tracking of lineages with CloneSweeper. bioRxiv 2026.01.30.700779 (2026).

22. Delley, C. L. & Abate, A. R. Modular barcode beads for microfluidic single cell genomics. Sci Rep 11, 10857 (2021).

23. Klein, A. M., Mazutis, L., Akartuna, I., Tallapragada, N., Veres, A., Li, V., Peshkin, L., Weitz, D. A. & Kirschner, M. W. Droplet barcoding for single-cell transcriptomics applied to embryonic stem cells. Cell 161, 1187–1201 (2015).

24. Clark, I. C., Fontanez, K. M., Meltzer, R. H., Xue, Y., Hayford, C., May-Zhang, A., D’Amato, C., Osman, A., Zhang, J. Q., Hettige, P., Ishibashi, J. S. A., Delley, C. L., Weisgerber, D. W., Replogle, J. M., Jost, M., Phong, K. T., Kennedy, V. E., Peretz, C. A. C., Kim, E. A., Song, S., Karlon, W., Weissman, J. S., Smith, C. C., Gartner, Z. J. & Abate, A. R. Microfluidics-free single-cell genomics with templated emulsification. Nat Biotechnol 41, 1557–1566 (2023).

25. Feng, Y., White, A. K., Hein, J. B., Appel, E. A. & Fordyce, P. M. MRBLES 2.0: High-throughput generation of chemically functionalized spectrally and magnetically encoded hydrogel beads using a simple single-layer microfluidic device. Microsyst Nanoeng 6, 109 (2020).

26. Karbaschi, M., Shahi, P. & Abate, A. R. Rapid, chemical-free breaking of microfluidic emulsions with a hand-held antistatic gun. Biomicrofluidics 11, 044107 (2017).

27. Stewart, M. P., Langer, R. & Jensen, K. F. Intracellular Delivery by Membrane Disruption: Mechanisms, Strategies, and Concepts. Chem Rev 118, 7409–7531 (2018).

28. Ruzzante, B., Fruzzetti, F., Cattaneo, M., Lauria Pinter, G., Marcuzzo, S., Candiani, G. & Bono, N. Harnessing osmotic shock for enhanced intracellular delivery of (nano)cargos. Int J Pharm 669, 125008 (2025).

29. Medepalli, K., Alphenaar, B. W., Keynton, R. S. & Sethu, P. A new technique for reversible permeabilization of live cells for intracellular delivery of quantum dots. Nanotechnology 24, 205101 (2013).

30. Paramasivam, P., Stöter, M., Corradi, E., Dalla Costa, I., Höijer, A., Bartesaghi, S., Sabirsh, A., Lindfors, L., Yanez Arteta, M., Nordberg, P., Andersson, S., Baudet, M.-L., Bickle, M. & Zerial, M. Quantitative intracellular retention of delivered RNAs through optimized cell fixation and immunostaining. RNA 28, 433–446 (2022).

31. Sudji, I. R., Subburaj, Y., Frenkel, N., García-Sáez, A. J. & Wink, M. Membrane Disintegration Caused by the Steroid Saponin Digitonin Is Related to the Presence of Cholesterol. Molecules 20, 20146–20160 (2015).

32. Zheng, X. & Gallot, G. Dynamics of Cell Membrane Permeabilization by Saponins Using Terahertz Attenuated Total Reflection. Biophys J 119, 749–755 (2020).

33. Adam, S. A., Sterne-Marr, R. & Gerace, L. Nuclear protein import using digitonin-permeabilized cells. Methods Enzymol 219, 97–110 (1992).

34. Cao, Y., Hjort, M., Chen, H., Birey, F., Leal-Ortiz, S. A., Han, C. M., Santiago, J. G., Paşca, S. P., Wu, J. C. & Melosh, N. A. Nondestructive nanostraw intracellular sampling for longitudinal cell monitoring. Proc Natl Acad Sci U S A 114, E1866–E1874 (2017).

35. Ilicic, T., Kim, J. K., Kolodziejczyk, A. A., Bagger, F. O., McCarthy, D. J., Marioni, J. C. & Teichmann, S. A. Classification of low quality cells from single-cell RNA-seq data. Genome Biol 17, 29 (2016).

36. Kasai, H., Allen, J. T., Mason, R. M., Kamimura, T. & Zhang, Z. TGF-beta1 induces human alveolar epithelial to mesenchymal cell transition (EMT). Respir Res 6, 56 (2005).

37. Keer, F. R., Al’Khafaji, A. M. & Blainey, P. C. Stitch-seq: Scalable CRISPR gene expression response profiling. bioRxiv 2026.04.27.719216 (2026).

38. Najafi, A., Jolly, M. K. & George, J. T. Population dynamics of EMT elucidates the timing and distribution of phenotypic intra-tumoral heterogeneity. iScience 26, 106964 (2023).

39. Cook, D. P. & Vanderhyden, B. C. Context specificity of the EMT transcriptional response. Nat Commun 11, 2142 (2020).

40. Tang, F., Lao, K. & Surani, M. A. Development and applications of single-cell transcriptome analysis. Nat Methods 8, S6–S11 (2011).

41. Wang, J., Rieder, S. A., Wu, J., Hayes, S., Halpin, R. A., de Los Reyes, M., Shrestha, Y., Kolbeck, R. & Raja, R. Evaluation of ultra-low input RNA sequencing for the study of human T cell transcriptome. Sci Rep 9, 8445 (2019).

