## Supplementary material for "Live Droplet Single-Cell RNA Sequencing": SI figures

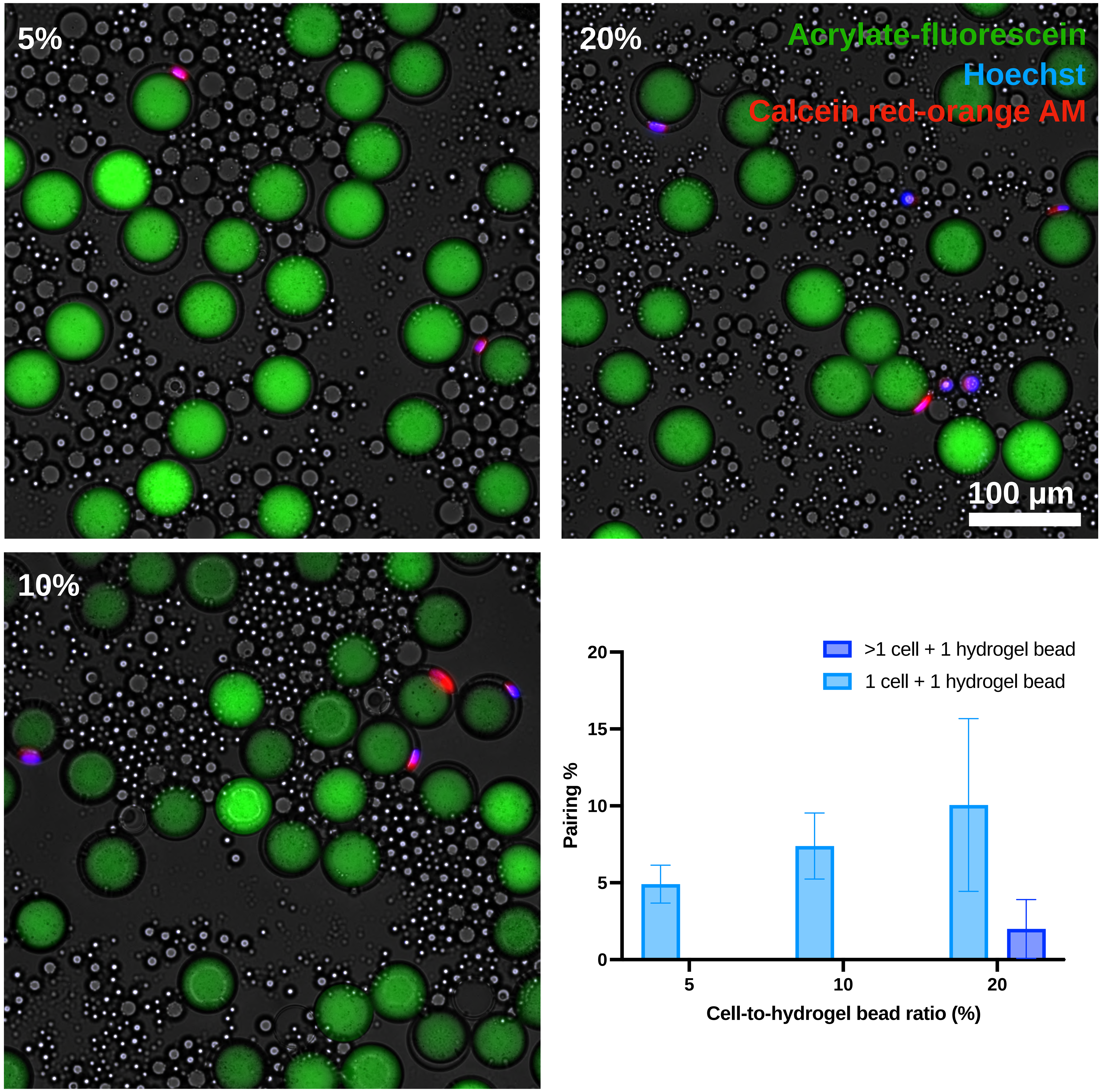


**Supplementary Figure 1: Single hydrogel bead and cell pairing (HEK293T).** Representative fluorescence images of droplets after particle-templated emulsification (PTE) at cell-to-hydrogel bead ratios of 5%, 10%, and 20%. Hydrogel beads are labeled with acrylate-fluorescein (green), cell nuclei with Hoechst (blue), and live cells with calcein red-orange AM (red). The bar graph shows the percentage of hydrogel beads paired with exactly one cell (light blue) or with more than one cell (dark blue) at each ratio. Data are shown as mean ± s.d. ($n$ = 5 fields of view). Scale bar: 100 µm.

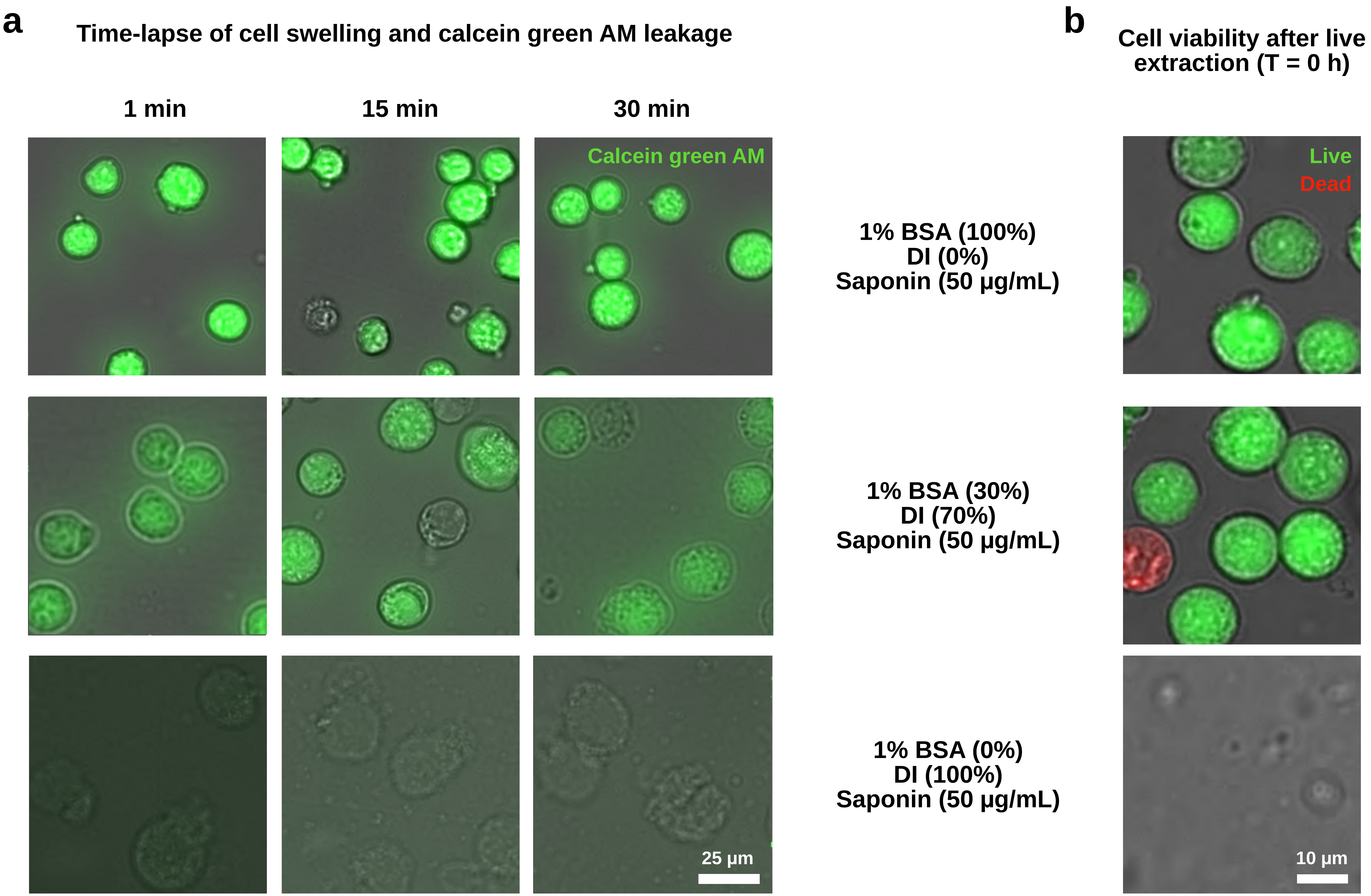


**Supplementary Figure 2: Time-course evaluation of extraction buffer treatment (HEK293T).** (**a**) Time-lapse images of calcein green AM-loaded HEK293T cells at 1, 15, and 30 min in extraction buffer containing 50 µg/mL saponin with 0%, 70%, or 100% (v/v) deionized (DI) water (balance 1% bovine serum albumin (BSA) in Dulbecco's phosphate-buffered saline (DPBS)). (**b**) Live/dead staining immediately after extraction (T = 0 h) under the same three buffer conditions. Live cells stained with calcein green AM (green) and dead cells stained with propidium iodide (red). Scale bars: 25 µm (**a**) and 10 µm (**b**).

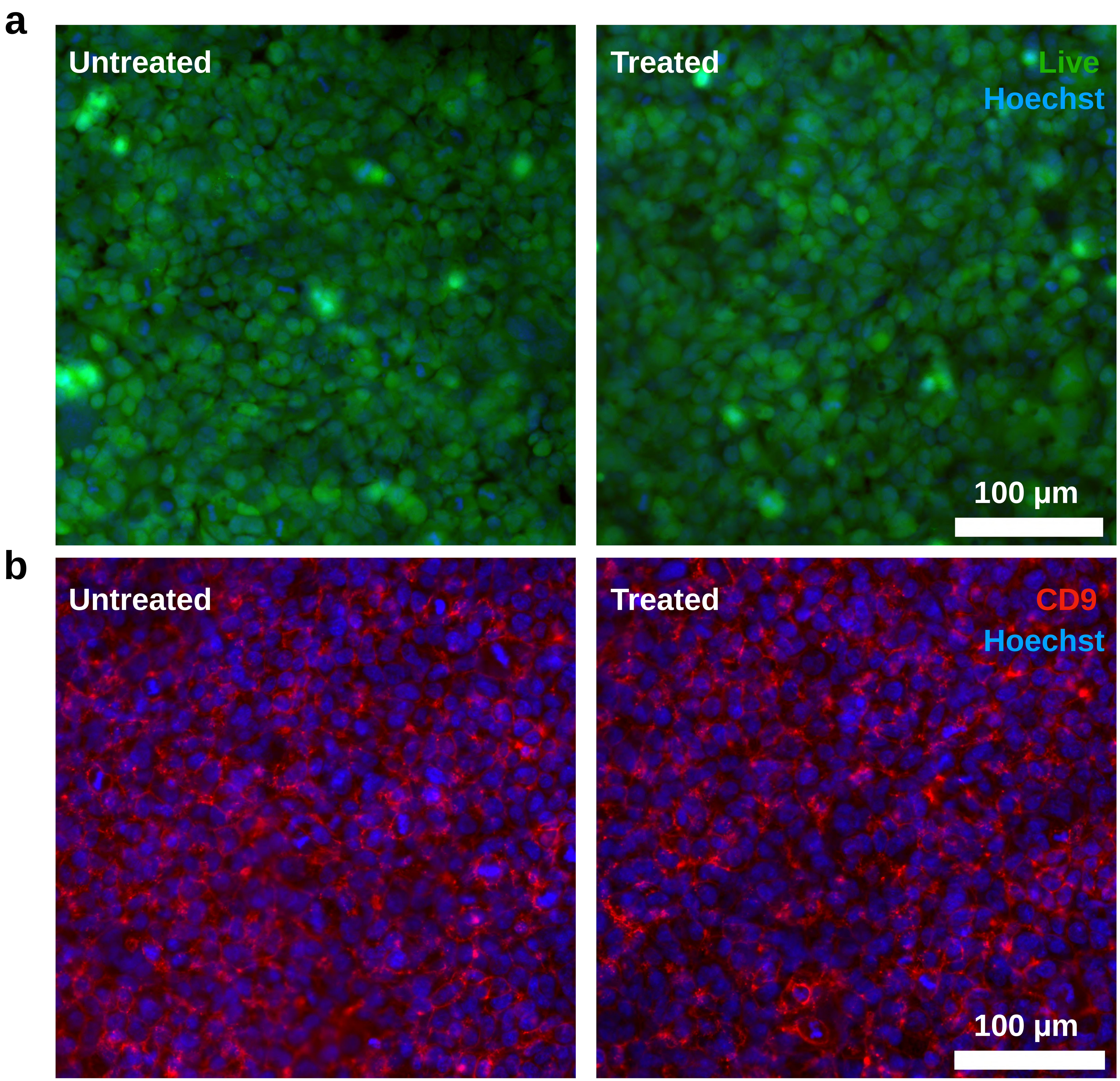


**Supplementary Figure 3: Cell viability and membrane validation (T = 24 h, HEK293T).** HEK293T cells were imaged 24 h after treatment with extraction buffer (70% (v/v) DI water, 50 μg/mL saponin) and compared with untreated controls. (**a**) Live cells stained with calcein green AM (green) and nuclei with Hoechst (blue). (**b**) Immunofluorescence staining of the membrane marker CD9 (red) with Hoechst nuclear counterstain (blue). Scale bars: 100 μm.

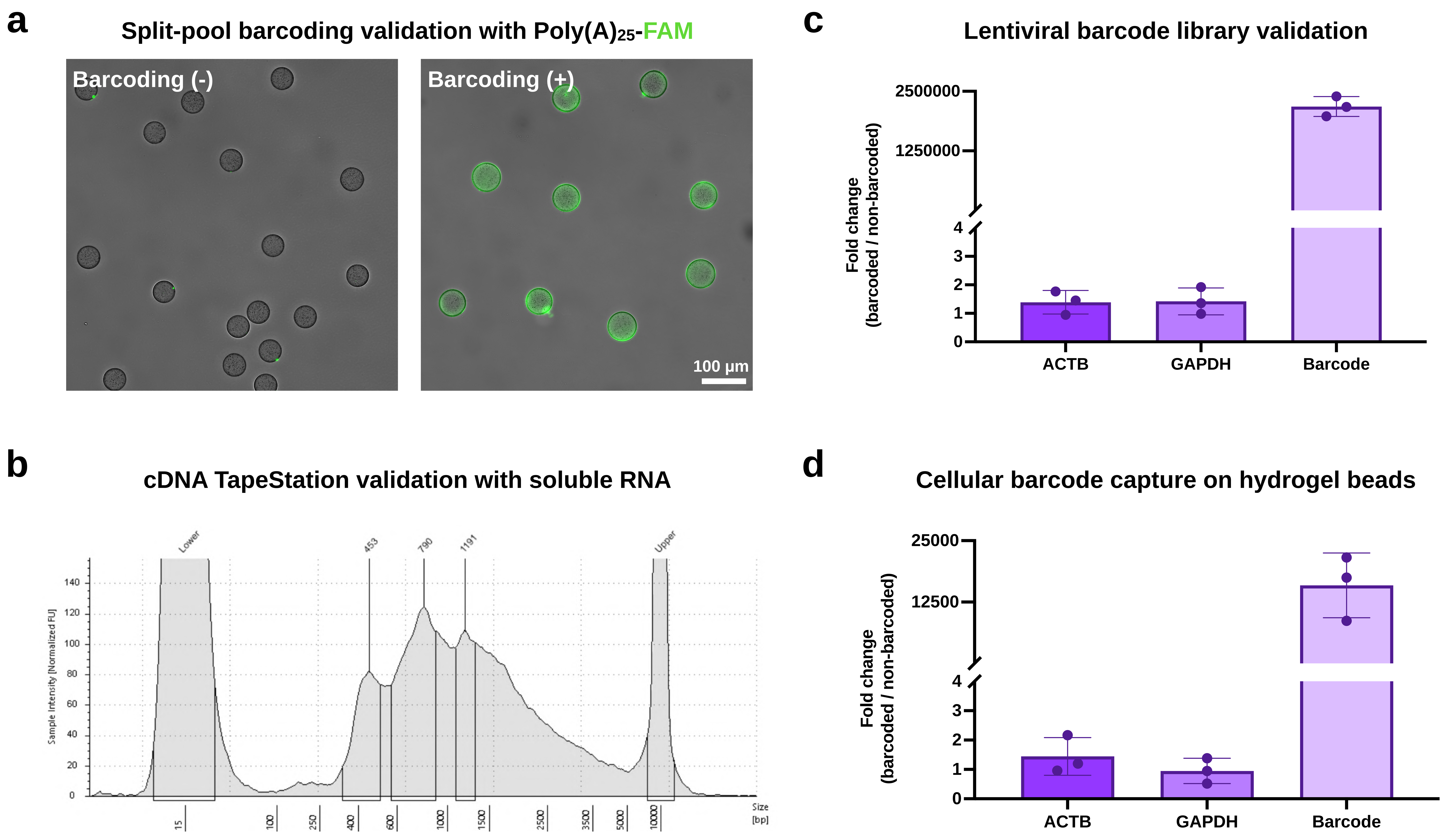


**Supplementary Figure 4: Hydrogel bead barcode validation and cellular barcode capture on hydrogel beads (A549).** (**a**) Fluorescence images of non-barcoded (−) and split-pool barcoded (+) magnetic hydrogel beads after hybridization with a FAM-labeled poly(A)$_{25}$ oligonucleotide. Scale bar: 100 µm. (**b**) TapeStation trace of cDNA generated from total RNA captured on barcoded hydrogel beads after reverse transcription, PCR amplification, and solid-phase reversible immobilization (SPRI) purification. (**c**) Reverse transcription quantitative PCR (RT-qPCR) of total RNA from barcoded versus non-barcoded A549 cells. (**d**) RT-qPCR of total RNA captured on barcoded hydrogel beads after co-incubation with total RNA from barcoded or non-barcoded A549 cells. In (**c**) and (**d**), the cellular barcode and the control genes *ACTB* and *GAPDH* are shown as fold change (barcoded / non-barcoded), normalized to *RPL13A*; y-axes are broken to show both ranges. Data are shown as mean ± s.d. ($n$ = 3 technical replicates).

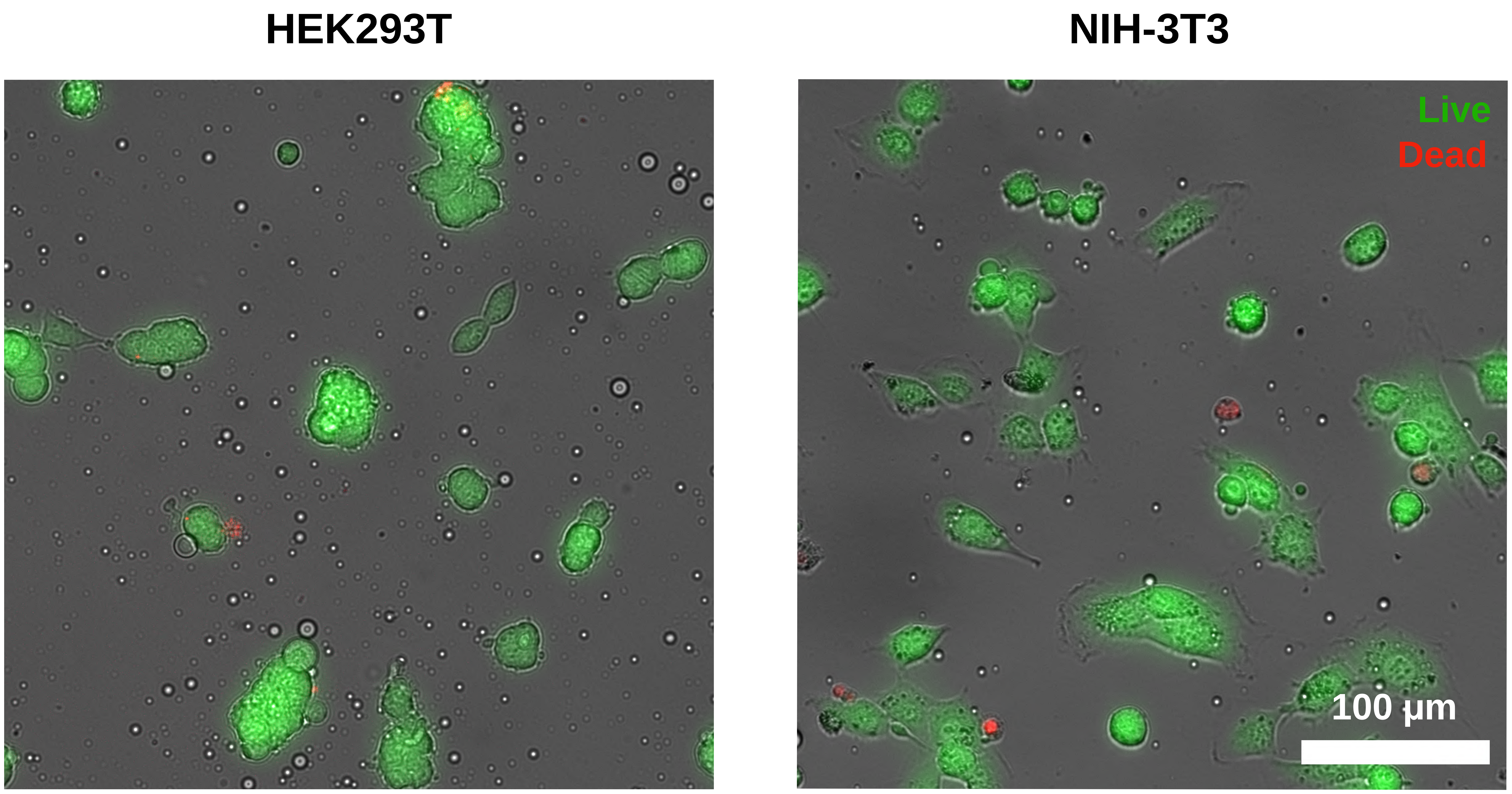


**Supplementary Figure 5: Cross-species cell viability (T = 24 h) post Live D-seq.** HEK293T (human) and NIH-3T3 (mouse) cells were processed separately by Live D-seq (70% (v/v) DI water, 50 µg/mL saponin) in parallel with the cross-species experiment, without mixing, so that each cell line could be identified. Cells were imaged 24 h after extraction. Live cells were stained with calcein green AM (green), and dead cells were stained with propidium iodide (red). Scale bar: 100 µm.

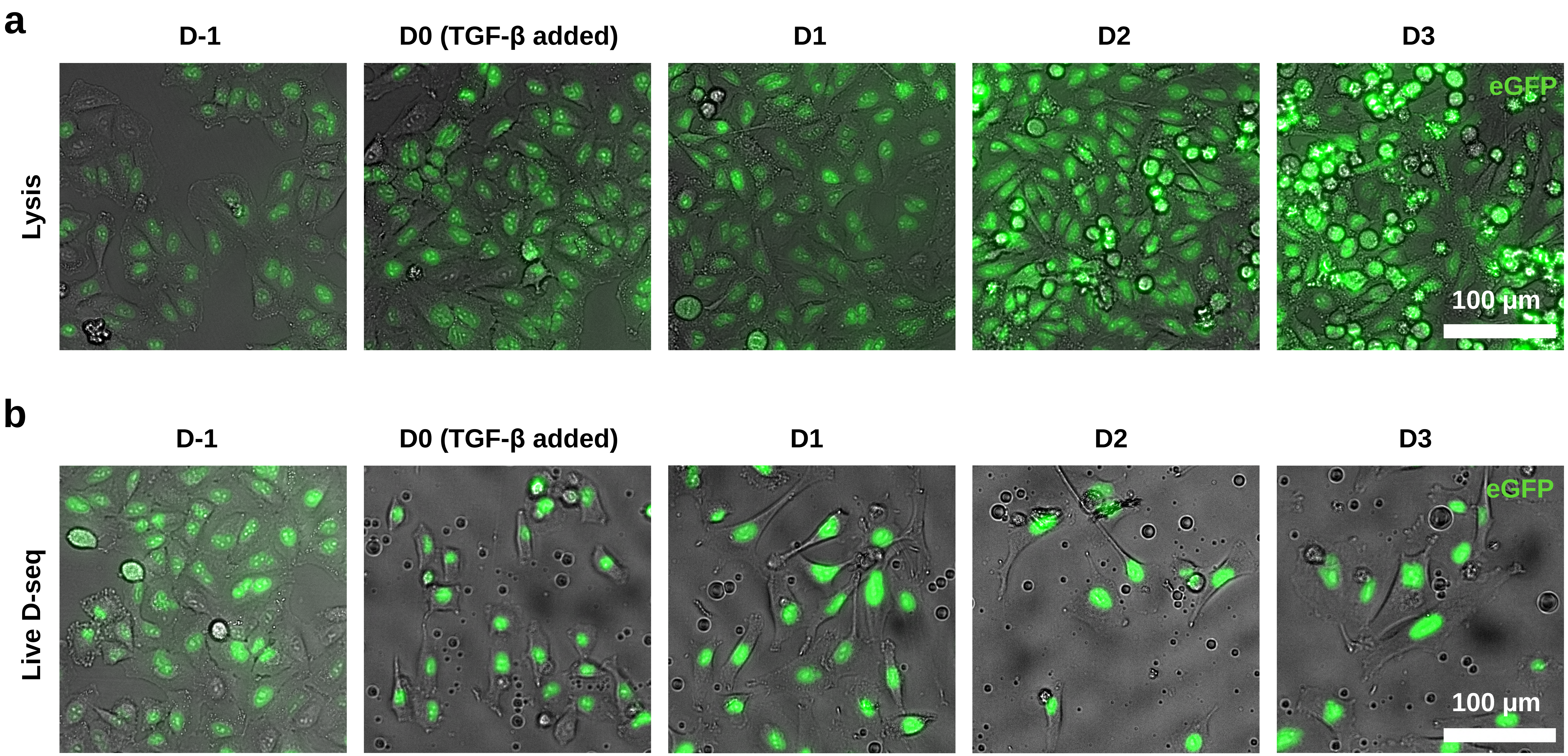


**Supplementary Figure 6: Longitudinal imaging of cells during EMT (A549).** Merged bright-field and enhanced green fluorescent protein (eGFP) fluorescence images of barcoded A549 cells from one day before (D-1) to three days after (D3) TGF-β addition. TGF-β (4 ng/mL) was added at D0. (**a**) Lysis controls: unsampled cultures that were lysed for single-cell RNA sequencing (scRNA-seq) at each time point. (**b**) Cells sampled by Live D-seq at D-1 and D1 and returned to culture between samplings. Scale bars: 100 µm.

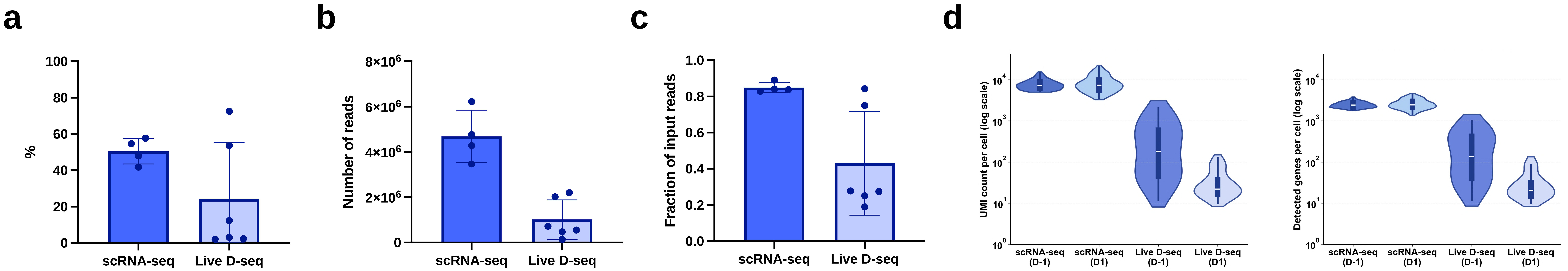


**Supplementary Figure 7: Sequencing metrics for the EMT longitudinal profiling experiment. Library metrics for scRNA-seq and Live D-seq.** (**a**) Sequencing saturation. (**b**) Number of sequencing reads. (**c**) Fraction of input reads uniquely mapped. (**d**) Unique molecular identifier (UMI) counts (left) and detected genes (right) per cell at D-1 and D1, shown on a log scale. Only cells that passed quality filtering and were used for downstream analyses (lineage tracing, principal component analysis (PCA), and gene expression comparisons) are shown. UMIs and genes were counted across all genes, without gene filtering. In (**a–c**), each point represents one library; data are shown as mean ± s.d.
